# Neuronal ROS-Induced Glial Lipid Droplet Formation is Altered by Loss of Alzheimer’s Disease-associated Genes

**DOI:** 10.1101/2021.03.03.433580

**Authors:** Matthew J. Moulton, Scott Barish, Isha Ralhan, Jinlan Chang, Lindsey D. Goodman, Jake G. Harland, Paul C. Marcogliese, Jan O. Johansson, Maria S. Ioannou, Hugo J. Bellen

## Abstract

A growing list of Alzheimer’s disease (AD) genetic risk factors is being identified, but the contribution of these genetic mutations to disease remains largely unknown. Accumulating data support a role of lipid dysregulation and excessive ROS in the etiology of AD. Here, we identified cell-specific roles for eight AD risk-associated genes in ROS-induced glial lipid droplet (LD) formation. We demonstrate that ROS-induced glial LD formation requires two ABCA transporters (*ABCA1* and *ABCA7*) in neurons, the APOE receptor (*LRP1*), endocytic genes (*PICALM*, *CD2AP*, and *AP2A2*) in glia, and retromer genes (*VPS26* and *VPS35*) in both neurons and glia. Moreover, ROS strongly enhances Aβ42-toxicity in flies and Aβ42-plaque formation in mice. Finally, an ABCA1-activating peptide restores glial LD formation in the APOE4-associated loss of LD. This study places AD risk factors in a neuron-to-glia lipid transfer pathway with a critical role in protecting neurons from ROS-induced toxicity.

## Introduction

Alzheimer’s disease (AD) is a neurodegenerative disorder characterized by memory loss and cognitive impairment (Rogaeva et al., 2006). AD affects ∼2% of the American population and defines ∼70% of dementia cases (Masdeu, 2020). AD is pathologically defined by the aberrant accumulation of Amyloid-β peptides (Aβ) into extracellular plaques and hyperphosphorylated-Tau into neurofibrillary tangles (NFT). Aβ-plaques primarily consist of Aβ42 fragments formed by the sequential cleavage of neuronally-expressed amyloid precursor protein (APP) (Behl, 1997; Karch and Goate, 2015; LaFerla and Green, 2012; Rogaeva et al., 2006). Tau, encoded by *MAPT*, is expressed in neurons, and is required for the assembly and stability of microtubules (MTs). Tau hyperphosphorylation inhibits its normal activity and drives its fibrillization into NFT (Mietelska-Porowska et al., 2014; Wang and Mandelkow, 2016). Both Aβ-plaques and NFT are neurotoxic in model organisms and the amyloid cascade hypothesis postulates that Aβ-plaque formation drives NFT formation (Bloom, 2014; Götz et al., 2007, 2011; Hardy et al., 2002). Thus, Aβ is a major focus in determining how AD is initiated and has been a target of many AD therapeutics (Bloom, 2014).

AD is currently classified as familial (FAD) or sporadic (SAD). FAD accounts for 2-3% of AD cases and is associated with autosomal dominant mutations in *APP*, *PSEN1*, or *PSEN2* (Giau et al., 2019; Zhu et al., 2015). It is hypothesized that SAD is caused by a combination of genetic and environmental risk factors (De Strooper and Karran, 2016). In fact, Aβ-plaques can form with age or due to trauma in the absence of cognitive impairment, supporting the hypothesis that a combination of genetic and environmental insults are required to induce disease (De Strooper and Karran, 2016). Of interest, multiple genome-wide association studies (GWAS) have been performed on post-mortem AD tissue, identifying over 40 potential genetic risk factors that are significantly associated with disease (Kunkle et al., 2019; Lambert et al., 2013). Interestingly, some of the SNPs identified in these studies are in or near genes that encode proteins involved in lipid regulation (e.g. TREM2, ABCA7) and clathrin-mediated endocytosis (e.g. *BIN1*, *CD2AP*, *AP2A2*, *PICALM*, and *RIN3*), but the consequences of disrupting these genes/pathways in specific cell types needs further examination. (Karch and Goate, 2015; Wellington, 2004). The well-established AD-risk allele, the ε4 allele of *Apolipoprotein E* (*APOE*), *APOE4*, is by far the highest genetic risk factor in these studies. *APOE4* is found in ∼40-60% of AD individuals, varying by geographic region, and has a highly significant association with age of AD onset (Karch and Goate, 2015; Ward et al., 2012). Homozygous carriers for *APOE4* are 8-12 times more likely to develop AD than non-carriers (Michaelson, 2014). In contrast, individuals carrying the ε2 allele of *APOE* (*APOE2*) have reduced risk of developing AD, supporting that it is neuroprotective against AD development (Conejero-Goldberg et al., 2014; Huang and Mahley, 2014; Huynh et al., 2017; Li et al., 2020). While APOE functions to mediate lipid transfer between cells, the APOE4 variant has reduced lipid transport capacity (Hatters et al., 2006; Liu et al., 2017).

In addition to genetic risk factors, environmental insults likely modulate severity and onset of disease, including cellular oxidative stress caused by accumulation of reactive oxygen species (ROS). ROS can oxidize and damage proteins, lipids, and nucleic acids (Butterfield and Mattson, 2020; Grimm and Eckert, 2017). When properly regulated, ROS can provide beneficial effects to the cell (Ristow and Schmeisser, 2011; Thapa and Carroll, 2017) but damagingly high levels of ROS can occur with age as cellular control mechanisms become depleted, such as decreased antioxidant enzyme expression (Grimm and Eckert, 2017). Recent studies have found evidence of ROS elevation, specifically lipid peroxidation byproducts, in post-mortem tissue from individuals with pre-AD diagnoses, including preclinical AD and mild cognitive impairment as well as in AD brains (Allan Butterfield and Boyd-Kimball, 2018; Berbée et al., 2017; Bradley-Whitman and Lovell, 2015; Bradley et al., 2012; Butterfield, 2020; Peña-Bautista et al., 2019; Zabel et al., 2018). As AD progresses, ROS production is likely exacerbated by Aβ42-mediated neurotoxicity (Allan Butterfield and Boyd-Kimball, 2018; Tönnies and Trushina, 2017) and persistent neuroinflammation (Bisht et al., 2018; Regen et al., 2017), thus creating a vicious cycle that can drive disease progression.

The complexity of AD pathogenesis and progression is further expanded by the observation that many AD risk genes are expressed in glia in addition to neurons, suggesting that disruptions of these genes may impact multiple cell types in the brain. There is increasing evidence for an important role of dysregulation of glial lipid metabolism in AD (Kunkle et al., 2019; Marioni et al., 2018; Di Paolo and Kim, 2011; Wong et al., 2017). TREM2 and the phospholipase, PLCG2, control lipid metabolism in microglia and may aid the transition of microglia to a disease-associated state (Andreone et al., 2020; Nugent et al., 2020). Additionally, lipid sensing by TREM2 and lipid transport by ABCA7 is linked to the clearance of Aβ by glial cells (Aikawa et al., 2018; Wang et al., 2015). Glia are also important in the secretion of APOE, which can bind Aβ and facilitate its clearance (Fan et al., 2009; Robert et al., 2017). Interestingly, Alois Alzheimer repeatedly described “adipose saccules” in glial cells of AD patients over a century ago (A. Alzheimer, 1911; Alzheimer, 1907; Stelzmann et al., 1995) but, the link between neurodegeneration and lipid droplet (LD) accumulation in glia wasn’t described until recently (Van Den Brink et al., 2018; Liu et al., 2015; Zhang and Liu, 2017). In flies, we showed that elevated levels of ROS induces the formation of glial LDs by transferring peroxidated lipids produced in neurons to glia in a process mediated by the apolipoprotein, Glial Lazarillo (Glaz; homolog to human APOD), whose function can be replaced by human APOE (Liu et al., 2017) (Figure 1). The transfer of lipids from cultured vertebrate neurons that are stressed and physically separated from glia in a dish has also been documented to be dependent on APOE (Ioannou et al., 2019a; Liu et al., 2017). Glial LDs provide some neuroprotective capacity when ROS levels are elevated (Van Den Brink et al., 2018; Liu et al., 2017). In the *Drosophila* visual system, pigment glia that surround photoreceptor neurons, transport lactate to neurons through monocarboxylate transporters. Lactate is converted to pyruvate and Acetyl-CoA in order to support the TCA cycle in mitochondria. However, defective mitochondria are unable to optimally utilize this energy source and simultaneously producing ROS, which activates JNK and SREBP transcription factors that drive lipid synthesis using Acetyl-CoA. Newly generated lipids become peroxidated in the presence of ROS and are subsequently transported intracellularly via fatty acid transport proteins (FatP) to a heretofore unknown lipid exporter (Figure 1). After lipid export from neurons, lipids bind to apolipoproteins which allow extracellular lipid transfer and paracrine cellular uptake of lipids. The nearby pigment glia express the apolipoprotein, *Glaz*, allowing for lipid transfer to glia, via an unknown mechanism, for eventual sequestration of peroxidated lipids in LDs (Figure 1). The production and transfer of lipids from neurons to glia is a highly dose dependent process and reducing many of the players involved in the process by 50%, including *Lactate dehydrogenase*, *Pyruvate dehydrogenase*, *fatp*, and *Glaz* causes a very significant reduction in the accumulation of LD in pigment glia (Liu et al., 2015, 2017). Moreover, *APOE2/3* can restore LD formation in a *Glaz* loss-of-function mutant background, but *APOE4* cannot restore this function, leading to accelerated neurodegeneration (Liu et al., 2017). Human *APOE4* has a limited lipid-binding ability, suggesting that lipid clearance may be critical in AD (Hatters et al., 2006). It has been suggested that if the lipid binding affinity of *APOE4* could be enhanced, neurodegeneration could be prevented or delayed. A pharmacological agonist of *ABCA1* has been shown to restore *APOE4* lipidation and ameliorates Aβ42/Tau pathologies in a mouse model of human APOE expression (Boehm-Cagan et al., 2016a; Hafiane et al., 2015). Thus, evidence is mounting that lipids are inextricably linked with pathogenic mechanisms in AD and other neurodegenerative diseases (Chung et al., 2020; Griendling et al., 2016; Lin et al., 2019; Reed, 2011).

**Figure 1.**
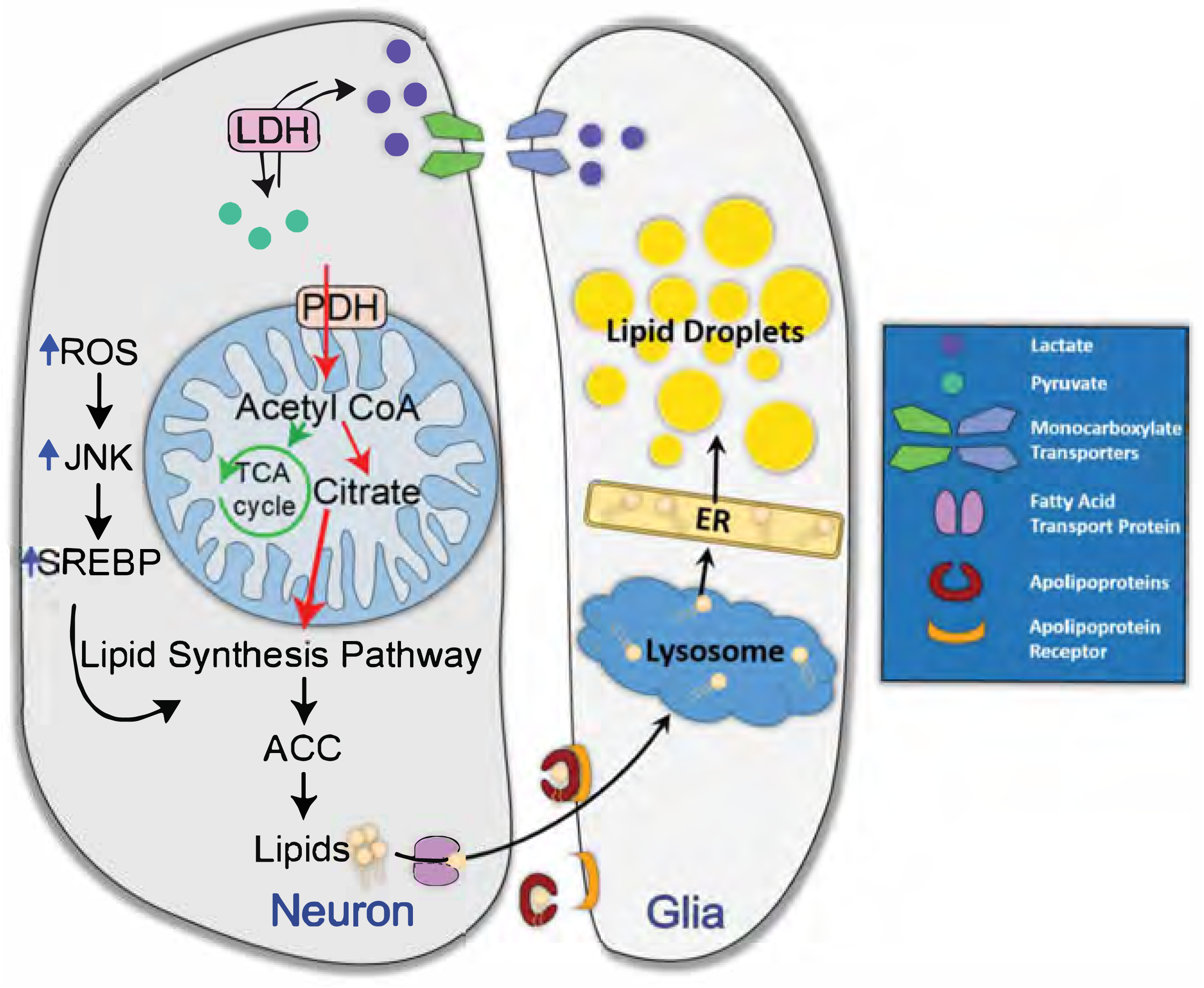
Neuron-to-glia lipid transfer for lipid droplet formation. Neurons and glia in the *Drosophila* visual system have a tightly regulated method of lipid production, transfer, and lipid droplet (LD) formation. Glia produce and transfer lactate through monocarboxylate transporters into neurons. Neurons convert lactate into pyruvate which feeds the TCA cycle in well-functioning mitochondrial. In defective mitochondria, ROS and citrate is produced which drives the synthesis of lipids by using Acetyl-CoA. Lipids are transported intracellularly via FatP and extracellularly via apolipoproteins where they are taken up in surrounding glia and incorporated into LDs. Adapted from Liu *et al*. (2017)

Here, we demonstrate that the fly homologs of multiple genes identified in AD GWAS and other clinical studies (*PICALM, CD2AP, AP2A2, ABCA1*, *ABCA7, LRP1*, *VPS26*, and *VPS35*) play a role in the formation of glial LDs, providing a mechanism by which these AD risk genes are involved in neuronal dysfunction. We demonstrate that two ABCA transporters are required in neurons, presumably for the export of lipids, and that six genes, involved in the uptake of APOE, are required in glia for the formation of glial LDs. Our data also show that elevated ROS synergizes with Aβ production to accelerate neuronal death in flies and exacerbates amyloid plaque formation in mice. Finally, we show that a peptide, previously shown to enhance the lipid-binding properties of *APOE4*, restores glial LD formation in a humanized fly model of *APOE4*. These data place AD-risk genes in a functional pathway that connects ROS with lipid production and Aβ toxicity, thus providing a possible therapeutic avenue for clinical intervention of AD based on utilization of antioxidants that can cross the blood-brain-barrier (BBB).

## Results

### ABCA transporters are required in neurons for glial LD formation

As we have previously shown that apolipoproteins are required for the transfer of peroxidated lipids from neurons to glia, we hypothesized that additional proteins are necessary for the export of lipids from neurons to the extracellular space where they lipidate apolipoproteins. Lipoproteins would then bind to a receptor on glial cells where endocytosis leads to the trafficking of the lipoproteins-rich endosomes to the lysosome for degradation of proteins and subsequent transport of the lipids to the ER for LD formation. We set out to assess the role of AD risk-associated proteins that export lipids from cells that are also expressed in neurons (Abuznait and Kaddoumi, 2012; Pereira et al., 2017).

The adenosine triphosphate-binding cassette transporter A1 (*ABCA1*) and A7 (*ABCA7)* encode lipid floppases that transfer lipids from the inner leaflet to the outer leaflet of the cell membrane making them available for export to APOE acceptor particles (Tarling et al., 2013; Turton and Morgan, 2013; Wahrle et al., 2004). Further, nonsense variants in both *ABCA1* and *ABCA7* have been associated with increased risk of developing AD (Bossche et al., 2016; Chang et al., 2019; Chen et al., 2016; Fehér et al., 2018; Teresa et al., 2020). *ABCA7* has also been identified as an AD risk locus in GWAS and is known to facilitate clearance of amyloid-β (Aikawa et al., 2018). Thus, the ABCA transporters are attractive candidates for neuronal export of lipids induced by ROS.

We set out to identify the fly homologs of ABCA1 and ABCA7 in the fly nervous system. The fly genome contains 10 putative ABCA transporters compared to 13 ABCA transporters in humans, and pairwise analysis of protein sequence does not reveal an obvious 1:1 homology. To identify fly ABCA proteins that share the greatest homology with ABCA1 and ABCA7 we assembled a gene tree of human and fly ABCA protein sequences. We found that fly genes *CG34120* and *Eato* grouped with human ABCA1/2/4/7/12/13 suggesting that these two fly genes may be orthologs of this group of human transporters (Figure 2A).

**Figure 2.**
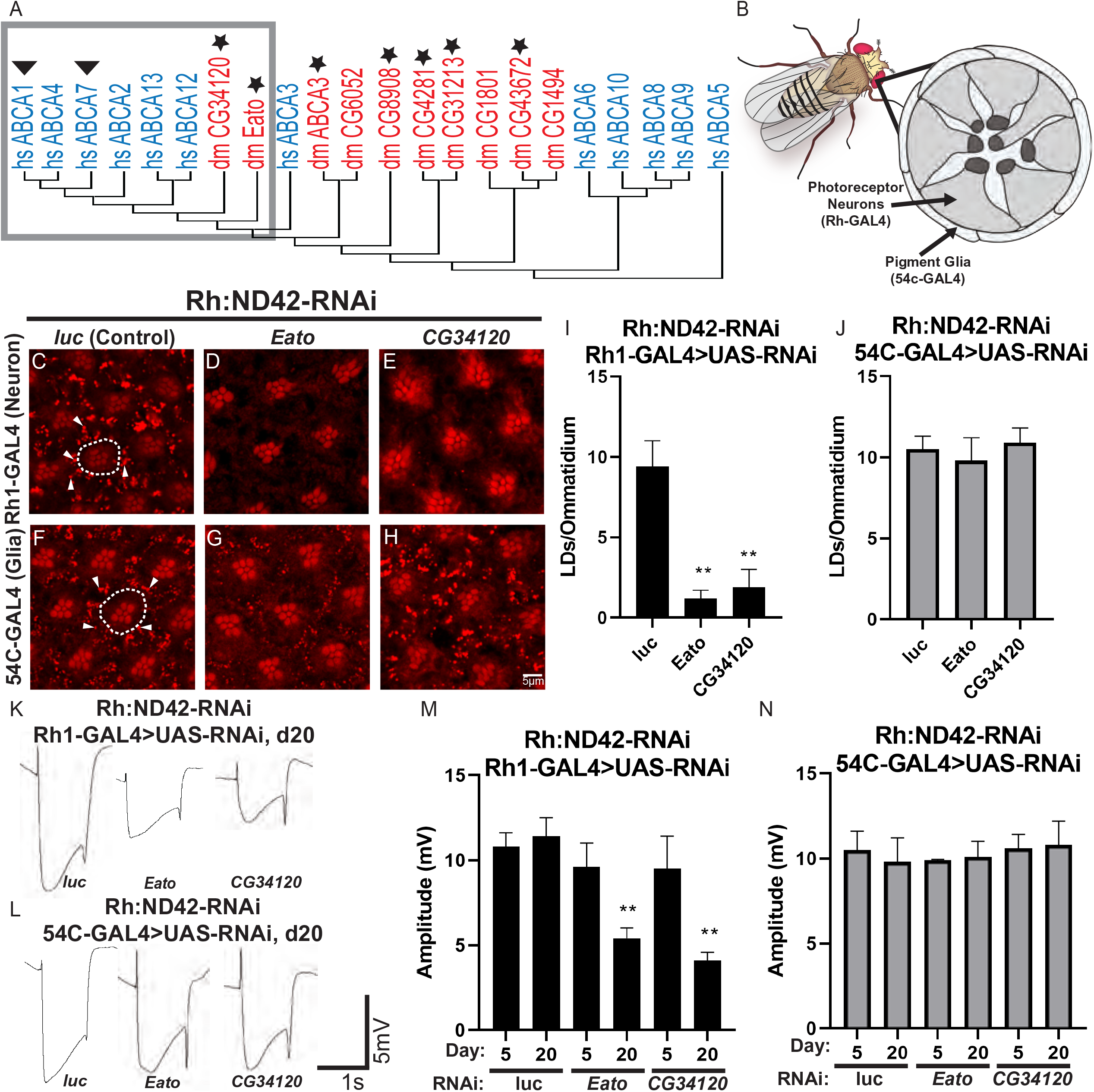
Two ABCA transporters are required in neurons for LD formation in glia. (A) Gene tree of fly (red) and human (blue) ABCA transporter protein sequences. Human ABCA1 and ABCA7 (triangles) have been implicated as AD risk genes. Stars indicate the fly genes that were assayed in this study. Grey box indicates monophyletic grouping of closest-related fly genes to AD risk genes and their paralogs. *Eato* and *CG34120* are the closest homologs to the risk genes *ABCA1* and *ABCA7*. (B) The *Drosophila* eye was utilized in this study as a model of lipid transfer between neurons and glia. There are 7 visible photoreceptor neurons (with central rhabdomeres) in each optical section through a fly ommatidium, which are surrounded by pigment glia. (C-J) Lipid droplet analysis in fly retina. To induce ROS specifically in photoreceptor neurons, an RNAi against *ND42*, a mitochondrial complex I subunit is expressed under the control of the Rhodopsin (Rh)-GAL4 driver. Animals are reared at 29°C under 12-hour light/dark conditions. ROS in neurons induces glial LD formation in control animals (C and F). The photoreceptor rhabdomeres stain positive with Nile red but photoreceptors (dashed lines) do not accumulate LD. In contrast, pigment glia accumulate LD (arrowheads). Knockdown of *Eato* and *CG34120* in neurons (D-E), but not in glia (G-H), suppress LD formation, quantified in (I-J), demonstrating a critical role for these genes in neurons for LD formation. (K-N) To assess the functional consequences of the lack of LD formation we performed electroretinograms (ERGs) at day 5 and day 20. Animals were housed at 29°C under 12-hour light/dark conditions, n≥10 animals per genotype. Representative ERG traces from animals with genotypes indicated. ERG amplitude quantification (K and L) show that glial knock down of either *Eato* or *CG34120* gene tested does not affect ERG amplitude. However, neuronal knock-down of these genes lead to a dramatic reduction of ERG amplitude over time, showing a progressive neurodegeneration. Hence, ROS induced LD formation in glia provide a protection against neurodegeneration.

To assess whether ABCA transporters function in glial lipid droplet formation in the presence of neuronal ROS we used RNAi mediated knockdown to reduce ABCA expression in glia and neurons. As in previous studies, we utilized a ROS-induced LD fly model in which ROS production is initiated in photoreceptor neurons via *Rh*-mediated expression of an RNAi targeting the mitochondrial complex I gene, *ND42* (Liu et al., 2015). This induces LD formation in surrounding pigment glia that can be visualized, after staining with Nile Red, via confocal microscopy (Figure 2B). In this background, we induced expression of RNAi targeting all the fly ABCA genes for which RNAi is effective, in either neurons, using *Rh-GAL4*, or glia, using *54C-GAL4* and stained for LD. We quantified the efficiency of RNAi-mediated knockdown of each ABCA gene targeted in our analysis. We found that RNAi constructs targeting *Eato*, *CG34120*, *CG8908*, *CG31213*, *CG1494*, and *ABCA3* all efficiently reduced the expression of their respective ABCA transporters (Figure S1A). We observed significantly reduced LD formation when *Eato* and *CG34120*, but not *CG8908*, *CG31213*, *CG1494*, or *ABCA3* were knocked down in photoreceptors with two independent RNAi lines (Figure 2C-E, data not shown) compared to a control RNAi (luciferase RNAi). In contrast, when these genes were downregulated in glia, no obvious reduction in LD formation was observed (Figure 2F-H). These data demonstrate that two fly ABCA transporters, *Eato* and *CG34120*, are required in photoreceptor neurons for glial LD formation.

Our previous data showed a neuroprotective role for glial LD formation under ROS (Liu et al., 2015, 2017). Hence, we assessed if loss of LD formation, caused by RNAi targeting *Eato* and *CG34120*, could lead to an age-dependent neurodegeneration. We tested whether neuronal or glial knockdown of *Eato* and *CG34120* leads to increased neuronal dysfunction, under ROS conditions induced by Rh:ND42 IR, using the electroretinogram (ERG) assay. ERGs serve as a functional readout of neuronal function and viability (Jaiswal et al., 2012). ERG amplitudes were quantified in young (5 days post eclosion) and aged (20 days post eclosion) flies expressing RNAi targeting *Eato, CG34120*, or a control gene, *Luciferase* (*luc*). We observed a significant reduction in ERG amplitude with age when *Eato* and *CG34120* were targeted by RNAi in neurons (Figure 2 K-L, quantified in M-N) arguing that these two ABCA transporters are required in neurons for the neuroprotective mechanism of glial sequestration of peroxidated lipids in LD.

In summary, these data support a role for *Eato* and *CG34120*, orthologs of the AD risk genes *ABCA1* and *ABCA7*, in lipid transfer. Both genes are required in neurons, but not glia, for glial LD formation, supporting that they mediate the export of peroxidated lipids from neurons to glia to protect neurons from the toxic effects of lipid peroxidation.

### The APOE receptor, LRP1, and the retromer are required for glial LD formation

Previous reports demonstrated that glial LD formation requires the fly apolipoprotein, Glaz, supporting the idea that once lipids are exported across the cell membrane of neurons, they bind to apolipoproteins in the extracellular space and are taken up by glia (Liu et al., 2017). Similar conclusions were derived from experiments with primary cultures of vertebrate neurons and glia showing a requirement for APOE for LD formation in glia (Ioannou et al., 2019b; Liu et al., 2017). In these experiments, the endocytosis inhibitor Pitstop2 completely blocked lipid transfer from neurons to glia. However it remained unclear whether the inhibition of endocytosis acts on neurons or glia. We therefore assessed the effects of reduced neuronal or glial expression of genes involved in receptor-mediated endocytosis on glial LD formation, beginning with the genes encoding previously defined apolipoprotein receptors, *LRP1* and *VLDLR* (fly LpR2) (Herz, 2009; Lane-Donovan and Herz, 2017; Rodríguez-Vázquez et al., 2015). RNAi targeting these genes were expressed in photoreceptor neurons (Rh-GAL4) or pigment glia (54C-GAL4), as before, to assess impacts on glial LD formation when ROS is induced in neurons. Loss of *LRP1* in glia, but not neurons, caused a significant reduction in LD formation (Figure 3A and E). In contrast, neither neuronal nor glial expression of *LpR2* RNAi, altered LD formation (Figure 3B and F). These data argue that lipids produced under neuronal ROS require *LRP1* in glia, specifically.

**Figure 3.**
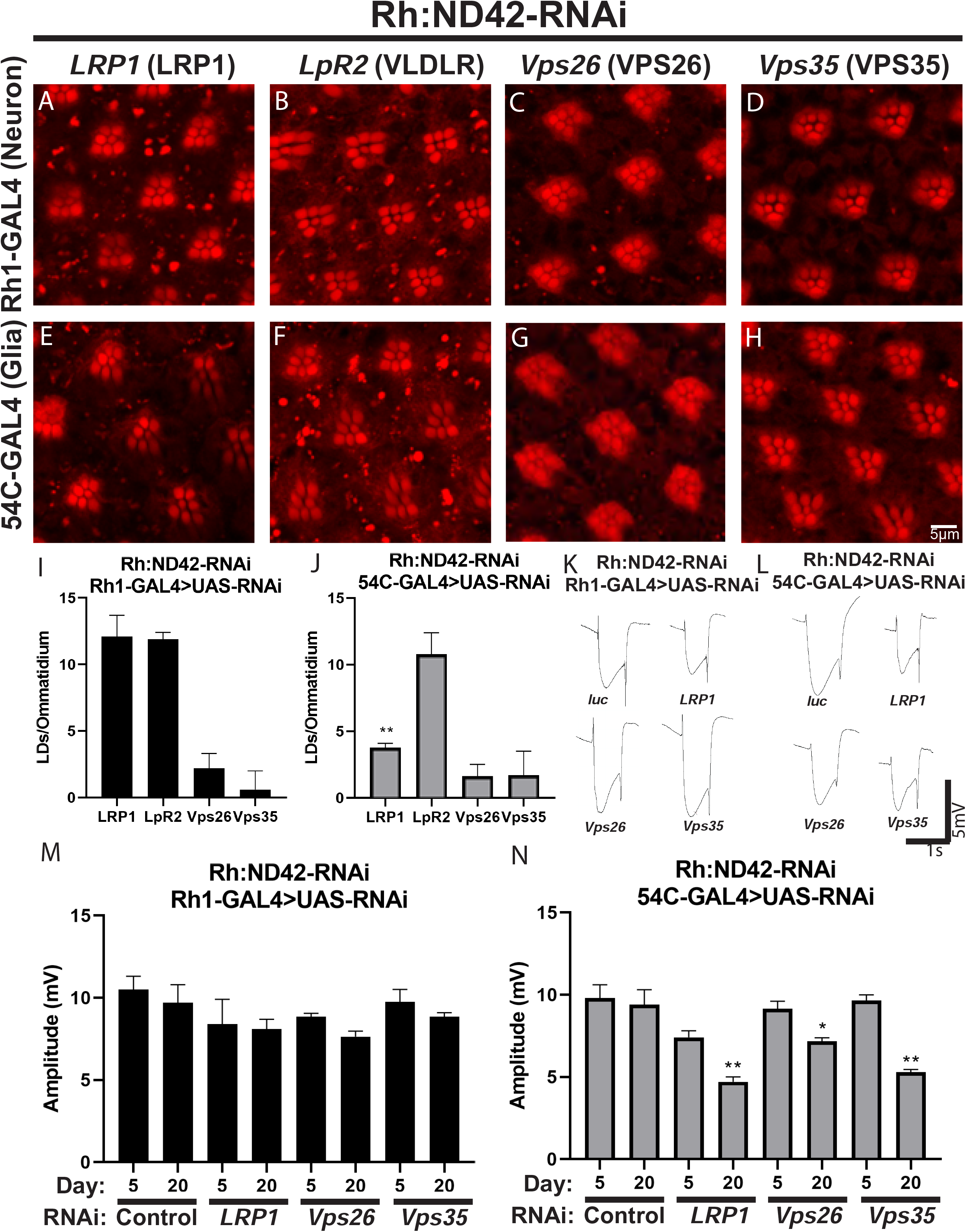
The APOE receptor, LRP1, and retromer components Vps26 and Vps35 are required for LD formation. (A-J) LD analysis in fly retina. ROS is induced in neurons and RNAi directed against the apolipoprotein receptors (*LRP1* and *LpR2*) or genes critical for retromer function (V*ps26* and *Vps35*) are expressed in neurons (Rh-Gal4, A-D) and pigment glia (54c-Gal4, E-H). Animals were reared at 29°C under 12-hour light/dark conditions. LRP1 is required in glia (E) but not in neurons (A) to form LD whereas LpR2 is not required in either cell (B, F). In contrast, the retromer proteins are required in both neurons and glia to form LD (C-D, G-H). LD number per ommatidium is quantified (I-J). (K-L) ERG assays were performed, as above, to assess neurodegeneration. Representative traces from animals with genotypes are shown (K-L). Quantification of ERG amplitude (M-N). Glial knockdown of *LRP1*, *VPS26*, or *VPS35* inhibits LD formation and is associated with an age-dependent neurodegeneration, consistent with a neuroprotective role of glial LD. In contrast, despite LD formation defects when *VPS26* or *VPS35* were knocked down in neurons, no or a mild neurodegeneration occurs suggesting ROS production or its effects are abrogated.

We then performed ERGs to assess if loss of *LRP1* in glia could impact age-dependent photoreceptor loss in animals with elevated levels of ROS in neurons. We found that, compared to control flies, glial, but not neuronal, knockdown of *LRP1* caused reduced ERG amplitudes in an age-dependent manner, indicative of photoreceptor degeneration (Figure 3K-N). These data suggest that the apolipoprotein receptor, *LRP1*, mediates glial import of peroxidated lipids produced in neurons and promotes glial LD formation and neuroprotection. These data also implicate receptor-mediated endocytosis as a critical pathway in glial LD formation. We therefore hypothesized that LRP1 recycling and vesicle endocytosis would similarly be important for LD formation.

The retromer serves critical functions in endocytosis, receptor recycling, and its loss in both photoreceptors and glia leads to neurodegeneration 20 days (Wang et al., 2014). Moreover, the retromer has been linked to several neurodegenerative diseases including AD (Berman et al., 2015; Muhammad et al., 2008) and deficiency of the retromer complex or its cargo proteins impairs endosomal trafficking of amyloid precursor protein (APP), resulting in the overproduction of β-amyloid (Aβ) (Qureshi et al., 2020; Zhang et al., 2018).

To determine the function of the retromer in LD formation, we targeted the retromer genes *VPS26* and *VPS35* with RNAi to knockdown their expression in our neuronal ROS model. *Vps26* and *Vps35* RNAi were expressed in neurons or glia and we assayed for LD formation and ERG amplitude. We found that loss of *Vps26* and *Vps35* in either neurons or glia leads to a significant reduction in glial LDs suggesting that the retromer is required in both neurons and glia for LD formation (Figure 3C-D and G-H). We assayed for age-dependent photoreceptor degeneration via ERG and found no worsening of photoreceptor function over time when *Vps26* or *Vps35* were knocked down in neurons (Figure 3K and M). In contrast, knockdown of *Vps26* and *Vps35* in glia caused an age-dependent reduction in amplitude indicative of neurodegeneration (Figure 3L and N). These data suggest that the neurodegeneration observed when glial retromer is lost may be caused by reduced uptake of peroxidated lipids in glia due to reduced apolipoprotein receptor recycling (Dhawan et al., 2020). The absence of ERG defects when expression of these genes is reduced in neurons at the time points assayed suggests that ROS production or the response to ROS production in neurons is blunted or delayed as the amplitude is decreased by 20 days, but is not statistically significantly different. However, the severe loss of ERGs documented in Wang *et al*. (2014) when either the Vps26 or Vps35 proteins is lost in both photoreceptors and glia suggest and additive or synergistic effect between neurons an glia and argues that the retromer is required in both cell types to maintain neuronal health.

### Endocytic AD-risk genes are required in glia for glial LD formation

A subset of AD-risk loci contain genes involved in endocytosis, including *BIN1*, *CD2AP*, *PICALM*, *AP2A2*, and *RIN3* (Van Acker et al., 2019; Nelson et al., 2020; Shen et al., 2020). This suggests that disruptions of this process may be important for AD pathogenesis. It is typically thought that these genes contribute to AD pathology through their well characterized function in synaptic transmission in neurons (Gan and Watanabe, 2018; Kaksonen and Roux, 2018; Seto et al., 2002; Takei and Haucke, 2001). However, we hypothesized that these endocytic genes also play a role in LD biogenesis by endocytosing lipids secreted from neurons for LD formation.

To examine a role for endocytic AD risk genes in LD formation, we examined LD formation and ERG phenotypes in animals in which homologs of AD-risk genes are targeted, via RNAi, in neurons and glia in the presence of *ND42* knockdown-mediated neuronal ROS induction. We found that knockdown of *cindr* (*CD2AP*), *Ap-2α* (*AP2A2*), and *lap* (*PICALM*) in glia, but not neurons, caused a reduction in LD formation (Figure 4A-D, G-J), thus implicating these genes as critical components of glial LD production. In contrast, reduced expression of *spri* (*RIN3*) and *amph* (*BIN1*) in neurons or glia did not affect LD production (Figure 4E-F, K-L). We observed an age-dependent decrease in ERG amplitude for the LD-critical genes *Ap-2α*, *lap*, and *cindr* when these were targeted in glia, suggesting that loss of these genes in glia promotes neurodegeneration due to failure to produce neuroprotective LDs in glia (Figure 4O-R). Taken together, these data suggest that loss or reduction of endocytosis in glia inhibits the neuroprotective effects of glial LD formation and implicates endocytosis in the process of neurodegeneration.

**Figure 4.**
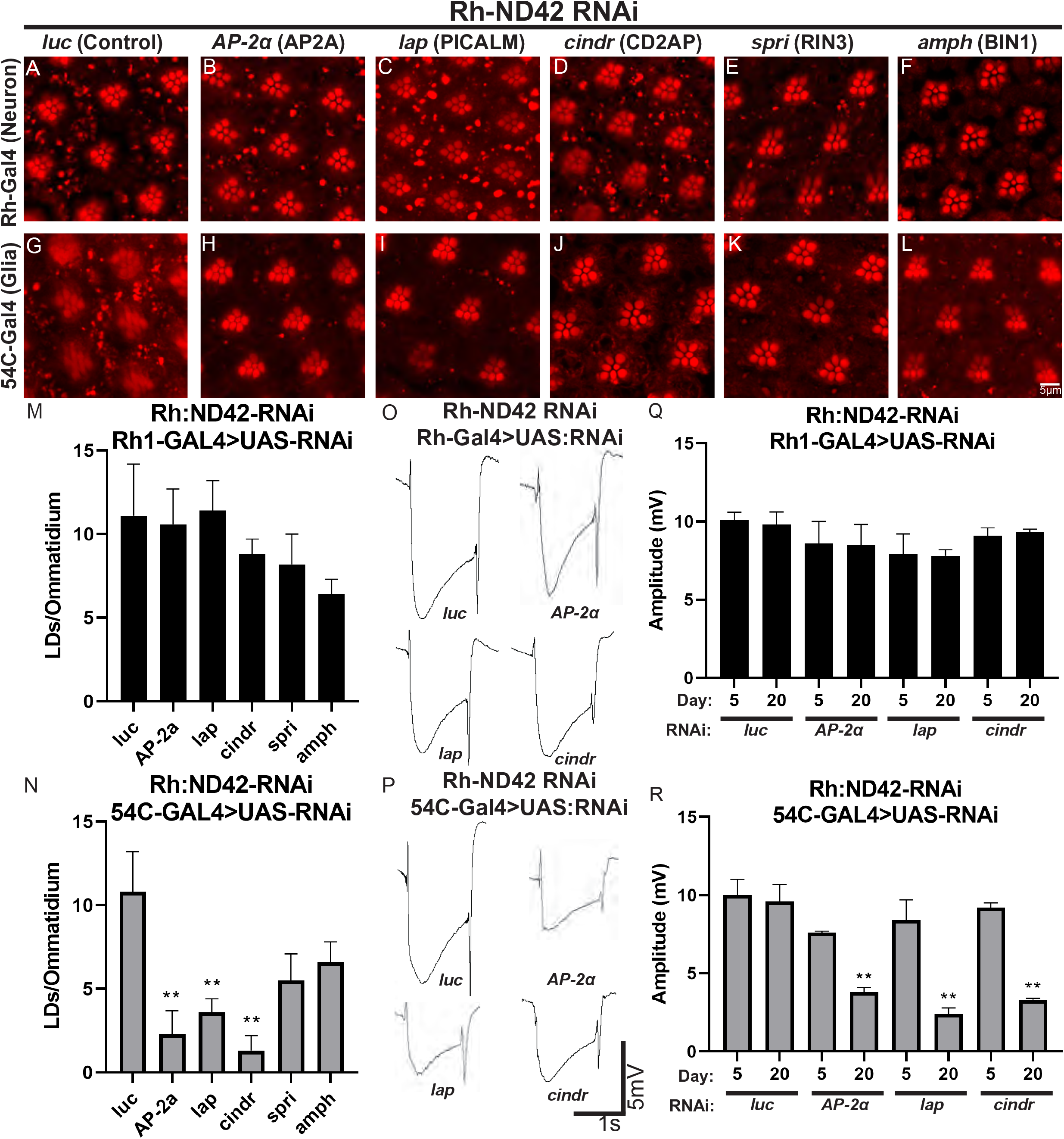
Alzheimer’s disease-associated GWAS genes are required in glia for LD formation upon neuronal ROS induction. (A-N) Lipid droplet analysis in fly retina. ROS is induced in neurons and RNAi directed against homologs of 5 GWAS genes in photoreceptor neurons (A-F) or glia (G-L). Animals are housed at 29°C under 12-hour light/dark conditions. Expression of RNAi against any genes tested in neurons do not affect the formation of LD in glia significantly (A-F). In contrast, RNAi targeting *AP-2a*, *lap*, and *cindr* (but not in a statistically significant manner in *spri* and *amph*) in glia reduced LD formation significantly (G-L) as quantified (M-N). (O-R) ERG assays were performed to assess neurodegeneration. Animals are housed at 29°C under 12-hour light/dark conditions, n≥10 animals per genotype. Representative traces (O-P) and amplitude quantification (Q-R) demonstrate that neuronal knock down of any gene tested does not affect ERG amplitude. In contrast, glial knock-down of the genes lead to a reduction in LD formation (*AP-2a*, *lap*, and *cindr*) led to a significant reduction of ERG amplitude over time, showing a progressive neurodegeneration. Hence, these genes are required in glia to take up peroxidated lipids and their loss promotes neurodegeneration.

We next investigated whether clathrin-mediated endocytosis is required for the uptake of neuron-derived fatty acids in a mammalian cell culture system. Since knockdown of *lap* in glia (Figure 4) caused a significant reduction in LD formation, we chose to knockdown the mammalian homologue *PICALM* in astrocytes and test the effects of fatty acid transport. We used lentivirus to deliver three independent shRNAs to reduce PICALM protein compared to a non-targeting shRNA control in cultured astrocytes (Figure 5A and B). We incubated neurons with a fluorescently labelled fatty acid analog Red-C12 overnight and then co-cultured the labeled neurons with transduced astrocytes on different coverslips separated by paraffin wax (Figure 5C) (Ioannou et al., 2019b, 2019a). We found a significant reduction in the transfer of fluorescently labelled fatty acids to astrocytes when PICALM levels are reduced (Figure 5D and E). Note that even relatively minor reductions of *PICALM* in astrocytes lead to reduced lipid accumulations. These data show that clathrin-mediated endocytosis is critical for the internalization of neuron-derived fatty acids in a mammalian culture system.

**Figure 5.**
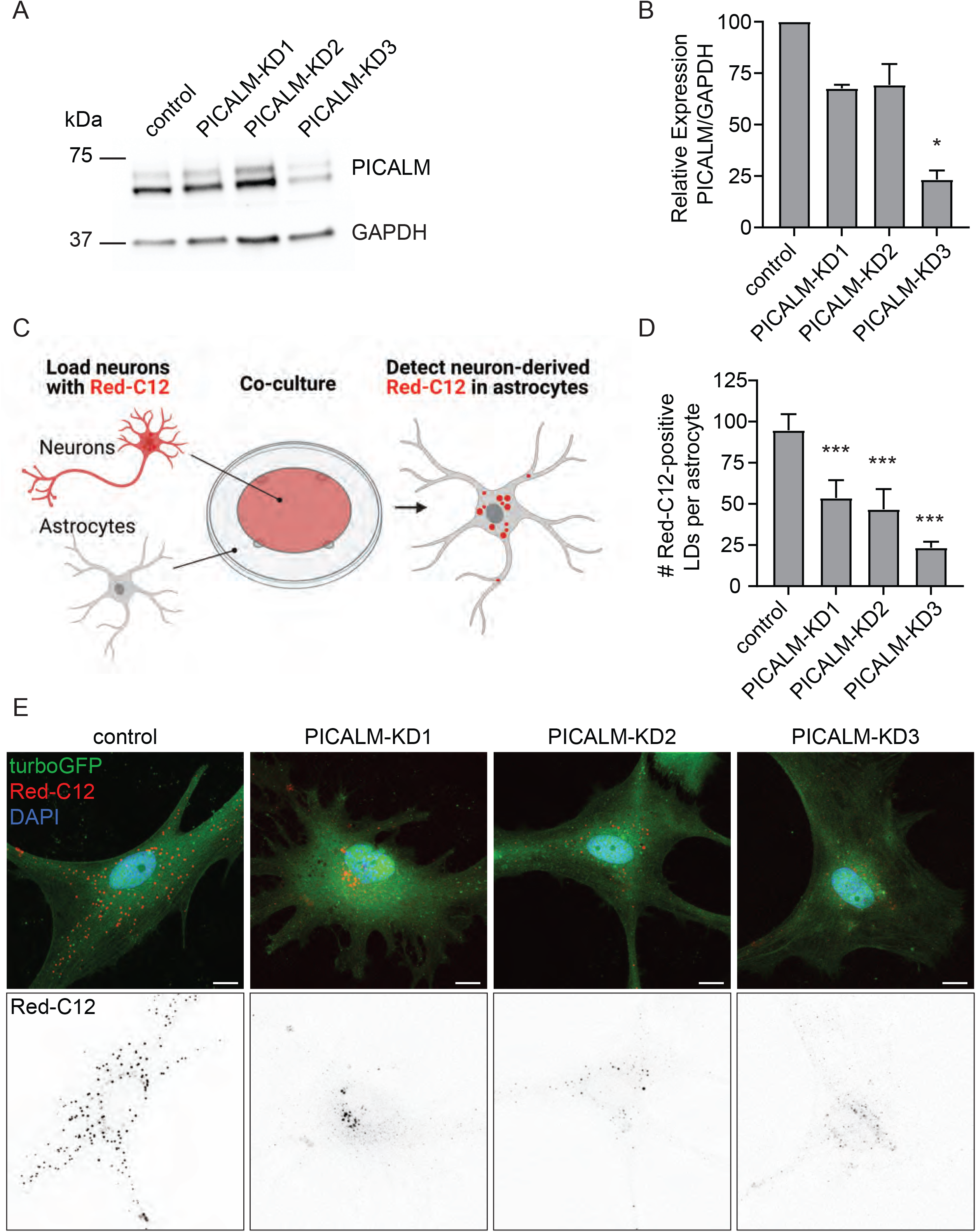
Lipid transfer between neurons and astrocytes is blunted by knockdown of PICALM. (A) Astrocytes were transduced with lentivirus expressing non-targeting shRNA (control), or three independent PICALM targeting shRNAs (KD1-3). Cell lysates were analyzed by Western blot for PICALM levels and GAPDH as a loading control. (B) Levels of PICALM from transduced astrocytes were quantified and normalized to GAPDH control. Mean ± SEM, Kruskal-Wallis test with Dunn’s posttest *p = 0.05 compared to control, n = 3 from three independent experiments. (C) Schematic of Red-C12 transfer assay. (D) Quantification of Red-C12-positive lipid droplets (LDs) in astrocytes. Mean ± SEM, One-way ANOVA with Dunnett’s posttest *** p < 0.001 compared to control, n = 6 from three independent experiments. (E) Representative maximum intensity projections of confocal images of transduced astrocytes following the assay. TurboGFP reporter expression marks transduced cells. Scale bars are 10 µm.

### Amyloid Beta synergizes with ROS in flies and mice

In AD, the plaque protein Amyloid β has lipophilic properties and has been shown to bind to the apolipoprotein receptor, LRP1, suggesting that altered lipid transfer may also alter amyloid deposition (Ermondi et al., 2015; Moreira et al., 2007; Verghese et al., 2013). There is also growing evidence that poorly lipidated APOE aggregates and acts as a seed for Aβ plaques (Lanfranco et al., 2020; Sharman et al., 2010; Verghese et al., 2013). This is supported by findings that ABCA1 loss-of-function leads to decreased APOE lipidation and increased amyloidogenesis (Koldamova et al., 2005; Wahrle et al., 2008). Further, as ROS-induced glial LD formation seems to be controlled by AD-associated risk genes and that peroxidated lipids accumulate in pre-AD patients (Allan Butterfield and Boyd-Kimball, 2018; Bradley-Whitman and Lovell, 2015; Bradley et al., 2012; Peña-Bautista et al., 2019), we hypothesized that ROS-induced lipid peroxidation may alter the effects of amyloid deposition in our model.

To test this hypothesis we expressed a secreted form of human Aβ42 in photoreceptor neurons, via *Rh-Gal4* (Chouhan et al., 2016). Low levels of ROS production, that avoids substantial neurotoxicity, was induced in animals by feeding them 25µM rotenone. Animals that expressed Aβ42 alone, or were fed rotenone alone, do not exhibit obvious signs of neuronal death in the fly retina and only a small number of glial LDs were observed in either condition (Figure 6A-B, D-E). In contrast, when Aβ42-expressing animals were fed 25µM of rotenone, by 10 days robust glial LD accumulation and a severe loss of rhabdomeres and overall PR morphology were observed (Figure 6F). These data demonstrate that extracellular Aβ42 strongly synergizes with low levels of ROS to induce premature neuronal death providing a mechanistic link between Aβ42 clearing and ROS.

**Figure 6.**
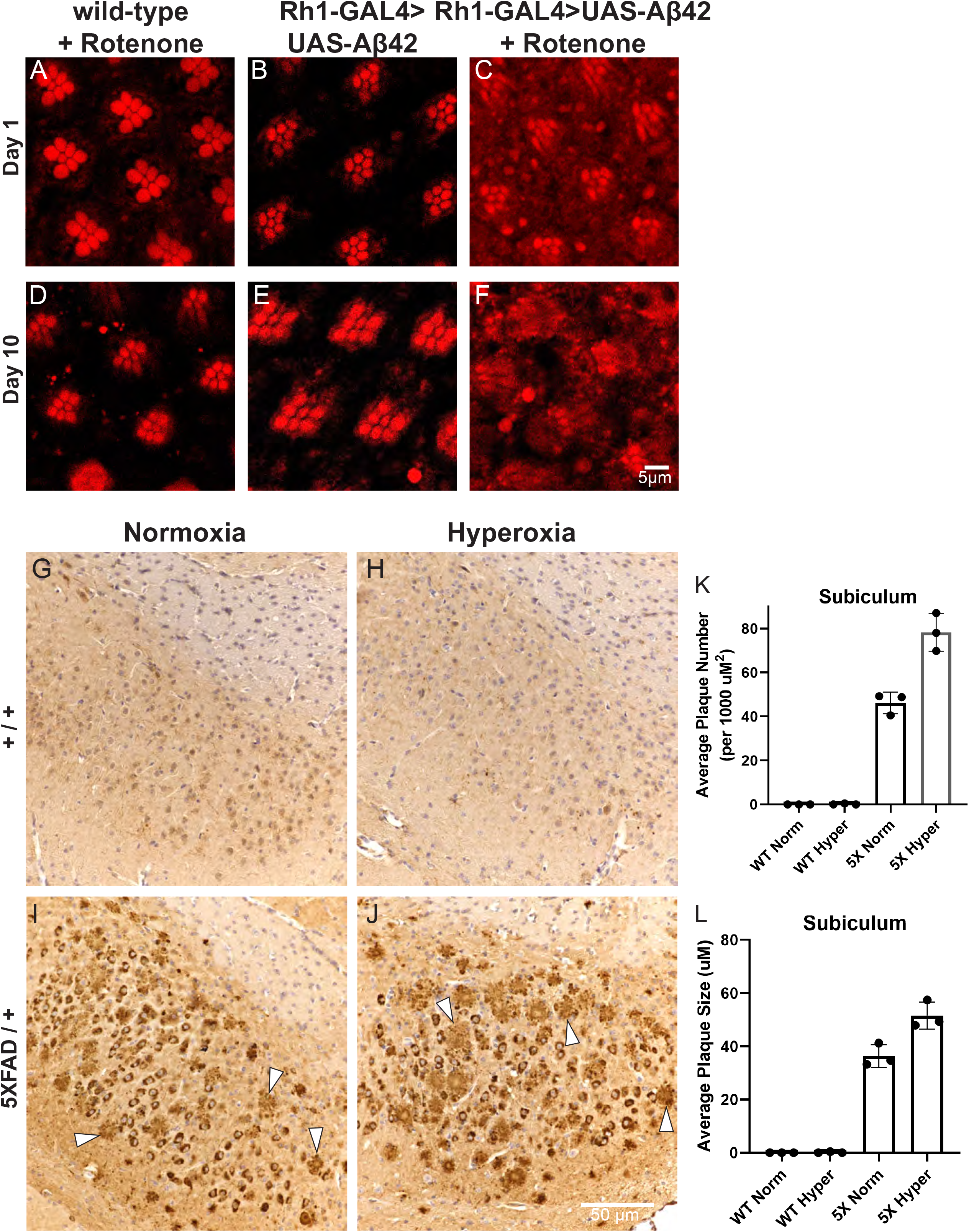
Elevated ROS and the presence of Aβ42 synergize to induce neurodegeneration in flies and mice. (A-F) Lipid droplet analysis in fly retina. Animals are housed at 29°C under 12-hour light/dark conditions with food changed daily; representative images of ≥10 animals per genotype. Wild-type flies exposed to 25 µM rotenone food at (A) 1 day or (D) 10 days post eclosion were compared with Aβ42-expressing flies at (B) 1 day post eclosion or (E) 10 days post eclosion and with Aβ42-expressing flies exposed to 25 µM rotenone food at (C) 1 day post eclosion and (F) 10 days post eclosion. Note the absence of LD formation with either treatment but the dramatic increase in diffuse Nile red staining and the demise of PR by day 10 showing that ROS and Aβ42 synergize to cause the demise of neurons. (G-L) Aβ42 immunohistochemical analysis of 4 mo. old mouse brain sections from wild-type mice reared in normoxic (G) or hyperoxic (H) conditions compared to 5XFAD mice reared in normoxic (I) or hyperoxic (J) conditions for 3 mos. prior to sacrifice. Arrowheads indicate plaques, n=3/genotype and treatment condition. Quantification of average plaque number (K) or plaque size (L) in the subiculum of mice from G-J is elevated in Aβ-expressing mice exposed to hyperoxia, showing that ROS induction enhances plaque formation.

We next tested for synergism between ROS and amyloid in a vertebrate model using the well-characterized 5XFAD mouse model (Jackson Labs) that expresses APP and forms Aβ-plaques by 4 months of age (Oakley et al., 2006). Previous studies have demonstrated that ROS can be induced in mice by rearing animals in a hyperoxic environment (Ferrari et al., 2017). We assembled cohorts of heterozygous 5XFAD mice and wild-type littermate controls and subjected them to either hyperoxia (55% O_2_) or normoxia (∼21% O_2_) conditions for 3 months beginning at the age of 4 weeks. Sagittal brain sections of the mice were probed for Aβ using established immunohistochemistry techniques (Sillitoe et al., 2008) (Figure 6G-J). We quantified plaque number and size in three regions of the brain, namely the cortex, hippocampus, and hindbrain. In each of these regions, plaque size and number observed in 5XFAD mice was significantly elevated in hyperoxia when compared to normoxia (Figure 6K-L, data not shown). Hence, ROS exacerbates Aβ-dependent phenotypes in mice similar to Aβ42 expressing flies.

### A pharmacological ABCA agonist peptide rescues APOE4 phenotypes

We previously reported that the AD-associated APOE4 allele was much less capable of mediating the transfer of lipids from neurons to glia (Liu et al., 2017). This work used APOE alleles to replace the fly apolipoprotein Glaz by inserting a T2A-GAL4 sequence into the *Glaz* Gene. This allele can drive the expression of any UAS transgene in the same spatiotemporal pattern as *Glaz* (Lee et al., 2018). Because we found that ABCA1 may be a critical lipid exporter in neurons exposed to ROS (see Fig. 2) and ABCA1 agonist peptides can promote APOE4 activity in AD mice (Boehm-Cagan et al., 2016a; Hafiane et al., 2015), we hypothesized that pharmacologically enhancing ABCA1 activity may restore LD formation in APOE4 flies.

We generated a fly line that expresses a genetically encoded version of the ABCA1 agonist peptide that was previously identified (Boehm-Cagan et al., 2016a, 2016b; Hafiane et al., 2015). The peptide sequence was cloned downstream of an Argos secretion signal to enable peptide release from the cell. Expression of the peptide was driven by the *Glaz^T2A-Gal4^* allele. We then elevated ROS levels in these animals by expressing an RNAi against *Marf*, the fly homolog of Mitofusin, under control of the *Rh* promoter, similar to Rh-ND42 IR as above (Liu et al., 2017). We confirmed the previous report that heterozygous *Glaz^T2A-Gal4^* animals have significantly reduced LD production (Liu et al., 2017) and showed that expression of the peptide does not alter LD formation in this background (Figure 7A and E). We next co-expressed the peptide with *APOE2, APOE3 or APOE4* alleles and compared it with expression of the *APOE* alleles alone (Figure 7B-D and F-J). Peptide expression with either *APOE2* or *APOE3* does not alter LD production in the absence of peptide expression. In contrast, expression of the peptide with *APOE4* restored LD formation, suggesting that the peptide can promote the lipidation of *APOE4* in flies and restore glial LD formation. These data show that this peptide indeed modifies the function of *APOE4* and restore its activity but has no impact on *APOE2* and *APOE3*.

**Figure 7.**
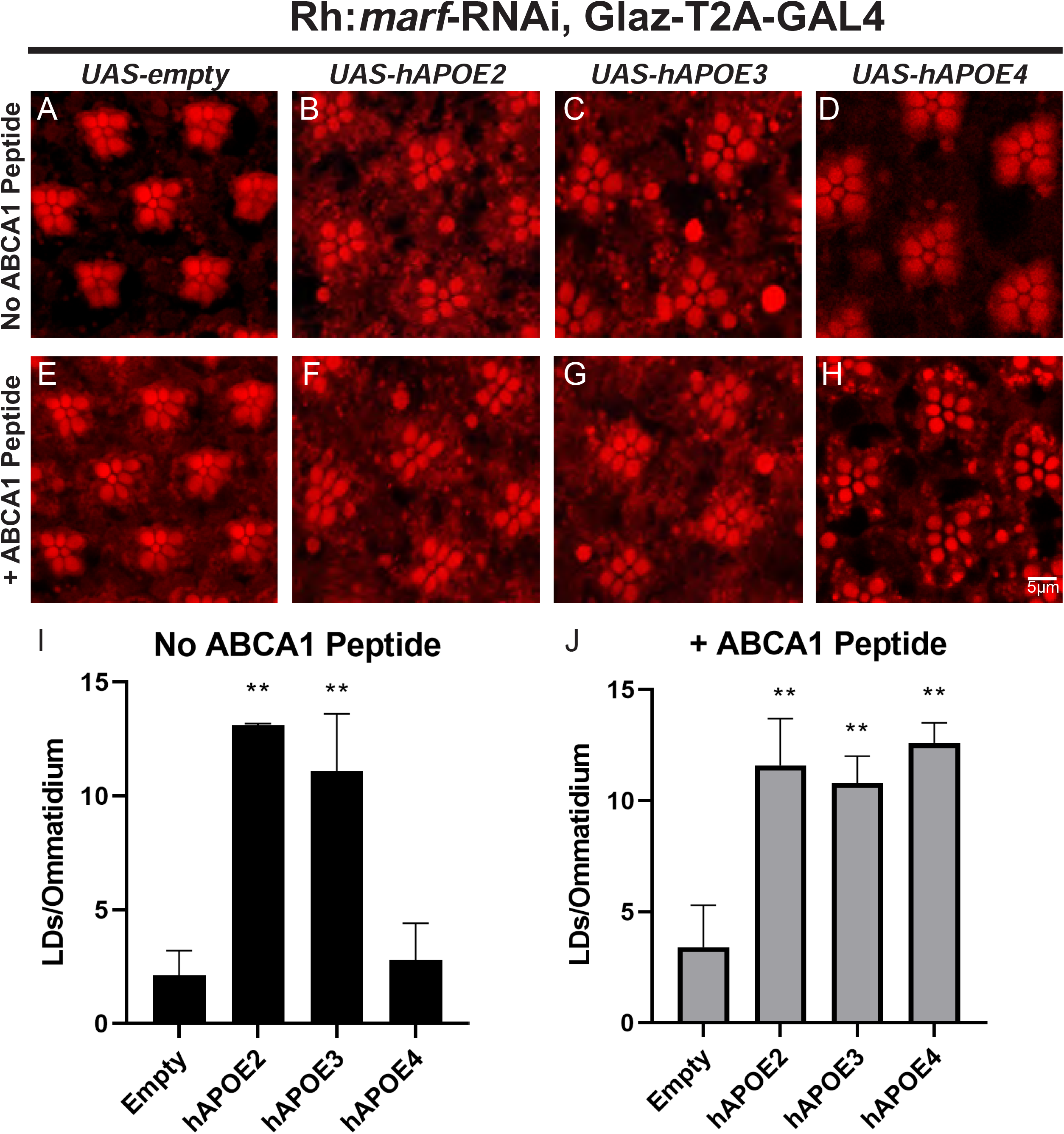
An ABCA1 agonist peptide rescues LD formation in the presence of APOE4. (A-J) LD analysis in fly retina. ROS was induced in photoreceptor neurons, as previously reported (Liu et al., 2015, 2017), using an RNAi against *marf*, the fly ortholog of mitofusin, under the control of Rh-GAL4. Animals are reared at 29°C under 12-hour light/dark conditions; representative images of ≥10 animals per genotype. We utilized a previously characterized allele of *Glial Lazarillo* (Glaz-T2A:Gal4). LD formation is inhibited in Glaz-T2A-Gal4/+ flies but can be restored by expressing human APOE2 or APOE3, but not APOE4. An ABCA1 agonist peptide was genetically encoded in the fly and expressed in the human APOE variant flies to assess LD formation. Expression of the peptide does not affect LD formation in the presence of APOE2 or APOE3, but fully restores LD formation in the APOE4 expressing flies (E-H) and quantified (I-J) showing that LD formation is strongly enhanced by this peptide.

## Discussion

We show that glia act to buffer against ROS produced in neurons by taking up peroxidated lipids and sequestering them in LDs (Figure 8). We found that glial sequestration of peroxidated lipids into LDs is neuroprotective and requires the function of genes involved in lipid export (*ABCA1* and *ABCA7*), lipid capture and transport (APOE) (Ioannou et al., 2019b; Liu et al., 2017), and receptor-mediated endocytosis (*LRP1*, *VPS26*, *VPS35*, *PICALM*, *CD2AP* and *AP2A2*). Notably, these genes have been implicated in AD and other neurodegenerative disorder risk genes (Hafiane et al., 2015; Kunkle et al., 2019; Lambert et al., 2013; Muhammad et al., 2008; Shinohara et al., 2017). These data suggest that AD risk could be elevated in the presence of partial loss-of-function variants of genetic risk factors by inducing lower efficiency of lipid transfer and peroxidated lipid sequestration in glial LD. This model predicts that the cumulative risk conferred by variants in the process of neuron-to-glia lipid transfer lies in the non-cell autonomous responses to neuronal ROS. Affected glia will be less well equipped to sequester peroxidated lipids, leading to increased levels of peroxidated lipids within and surrounding neurons, thus exacerbating neuronal demise. Although glia are well-equipped to sequester peroxidated lipids, they have a limited capacity to do so. Glia eventually succumb to the adverse effects of peroxidated lipid storage which eventually lead to a subsequent loss of neurons. It has been well documented that ROS levels are elevated with age and in multiple neurodegenerative diseases, including AD, and may be an important underlying cause of disease-associated neurodegeneration (Singh et al., 2019). Neurons have limited antioxidant capacity and innate mechanisms to respond to increased ROS by activating cellular responses (Burnside and Hardingham, 2017). Developing a better understanding of how protection against ROS is carried out should inform us about new ways to protect the nervous system from oxidative insult. Furthermore, understanding how these responses go awry may reveal ways to exogenously potentiate the antioxidant response. The identification of multiple genetic risk factors for AD that mediate glial LD formation (Figures 2-4), in combination with the observations that Aβ may be exacerbated by low levels of ROS (Figure 6) suggest that ROS-induced neuronal peroxidated lipid production and transfer to glia constitutes an important facet of the antioxidant toolkit (Liu et al., 2015, 2017). These data are also consistent with the previous observation that astrocytes are markedly more robust in handling and detoxifying ROS (Burnside and Hardingham, 2017). Hence, we argue that there is a critical need to reduce ROS in neurons in aging and the context of disease. We previously found that the fatty acid transporter protein, Fatp, is required in LD formation (Liu et al., 2017). However, it is not known to export lipids across membranes. Instead, Fatp may interact with lipid transport proteins such as ABCA transporters, known to transport lipids including cholesterol, phospholipids, fatty acids as well as other lipids across membranes (Neumann et al., 2017; Tarling et al., 2013). ABCA7, has been demonstrated to assemble high-density lipoprotein particles (Abe-Dohmae et al., 2004; Hayashi et al., 2005) and implicated in AD pathogenesis, first in an Icelandic population (Steinberg et al., 2015) and, later, in a larger AD cohort (Kunkle et al., 2019). The ABCA1 N1800H mutation, which has a low prevalence, is increased in frequency in the AD population and is associated with hemorrhagic stroke, consistent with clinical presentations in *APOE4* carriers (Nordestgaard et al., 2015). Interestingly, the sequence of the ABCA1 agonist CS6253 restored LD formation in *APOE4* flies but did not affect APOE2 or APOE3 function (Figure 7), supporting the findings of AD prevention in APOE4 targeted replacement mice (Boehm-Cagan et al., 2016a). This, together with our findings that ABCA transporters in the fly are required in neurons for glial LD formation (likely by mediating the export of peroxidated lipids) suggest a critical role for ABCA genes in proper lipid regulation in disease.

**Figure 8.**
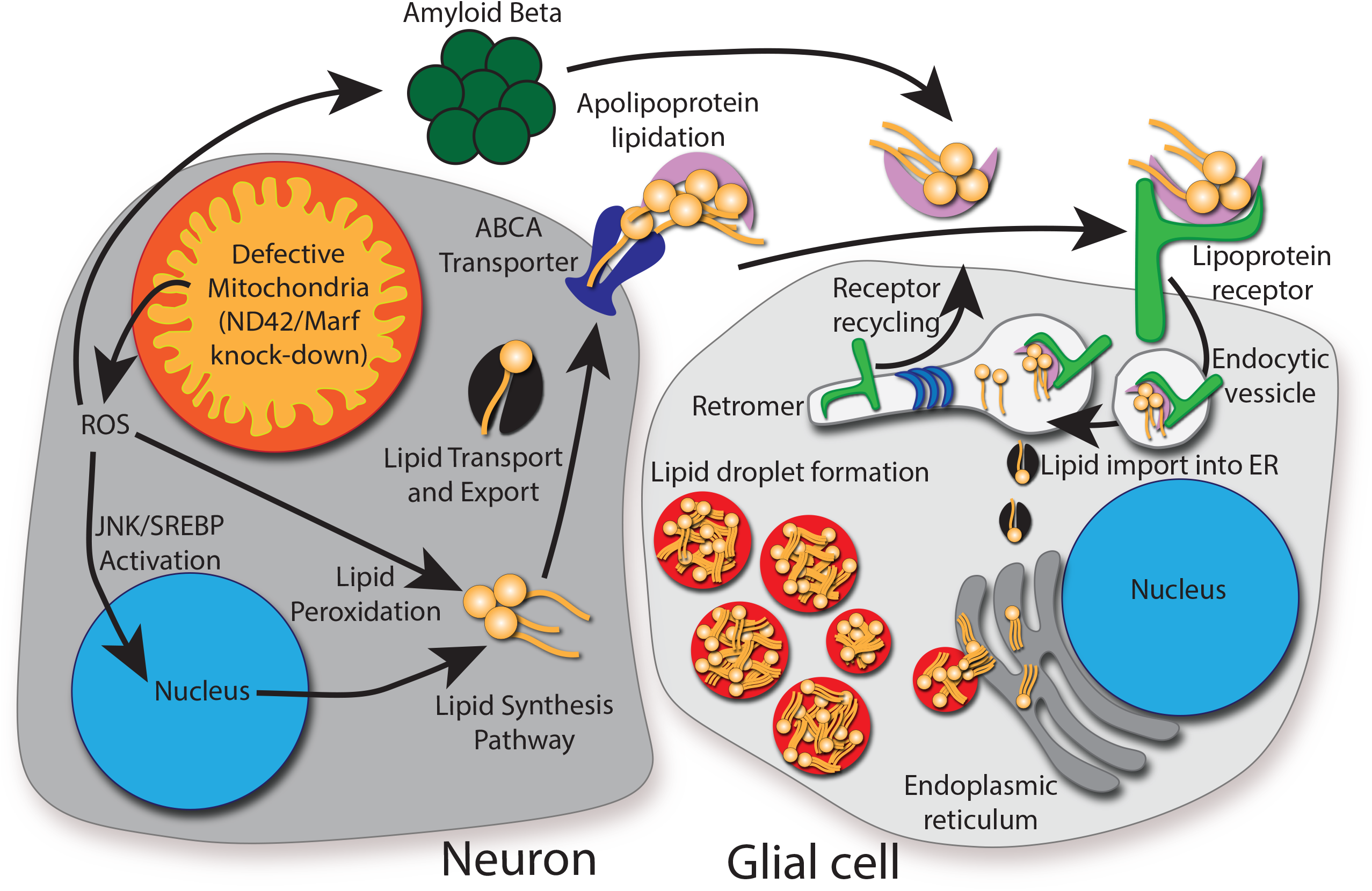
Model of lipid droplet accumulation and players identified in this study. We propose a model in which genetic (loss of ABCA, endocytic, or retromer genes) together with environmental insults (ROS) sensitize neurons to the presence of Amyloid accumulation to induce neurodegeneration. It is likely that this synergy between multiple insults severely exacerbates neuronal loss in disease. We demonstrated that lipid transfer between neurons and glia requires neuronal ABCA transporters, glial apolipoprotein receptors, and the retromer, which is required for LRP1 recycling. We propose that endocytosis of lipid particles are processed through lysosomes upon endocytosis. Lysosomes degrade Aβ42 and the lipids are shuttled to the ER to form LD. Hence, this transport of peroxidated lipids and Aβ42 provide dual protective effects.

The genes implicated in endocytosis studied in this work have often been studied in the context of synaptic transmission and/or neurodegenerative disease. *BIN1* is a membrane fission protein that regulates endocytic vesicle size in vertebrates, but it has been implicated in APP processing as well as Tau degradation (Van Acker et al., 2019; Ramjaun et al., 1997; Takeda et al., 2018). The fly homolog, *Amphiphysin* (*Amph*), regulates transverse tubule formation in muscles, which was also shown to be affected in vertebrate mutants (Lee et al., 2002) but *Amph* has not been implicated in endocytosis in flies to our knowledge (Razzaq et al., 2001; Zelhof et al., 2001). In contrast, *CD2AP* is a scaffolding protein that has been implicated in endocytosis and vesicle trafficking as well as APP sorting and processing in vertebrates (Furusawa et al., 2019; Harrison et al., 2016; Ubelmann et al., 2017). However, severe loss of function alleles of the fly homolog, c*indr*, affects synapse maturation as well as synaptic vesicle recycling and release (Ojelade et al., 2019). *PICALM* is a clathrin assembly protein that has been implicated in the import of γ-secretase and APP processing as well as Tau buildup (Van Acker et al., 2019; Baig et al., 2010). The fly homolog, *like-AP180* (*lap*) acts as a clathrin adaptor to promote clathrin-coated vesicle formation and restrict coated vesicle size as well as the efficacy of synaptic vesicle protein retrieval (Zhang et al., 1998). AP2A2, a member of the AP-2 adaptor protein complex, which aids in assembling endocytic components in flies and vertebrates, has implications in AD risk (González-Gaitán and Jäckle, 1997; Nelson et al., 2020). Finally, *RIN3* is a Rab5 guanine nucleotide exchange factor important for recruiting CD2AP and BIN1 to endosomes and has been implicated in APP accumulation and Tau phosphorylation regulation (Shen et al., 2020). Based on our data, three of these genes (*CD2AP*, *AP2A2*, and *PICALM*) play critical functions in glia for LD formation (Figure 4). Historically, because many of the endocytic AD-risk genes are known to play a critical role in synaptic transmission, it is thought that their role in AD pathology may occur due to loss of function of these genes in neurons. However, single cell RNA sequencing databases provide evidence for enriched expression of many of the endocytic AD risk genes in mouse/human glia, including those targeted in this study (Zhang et al., 2014, 2016) and our data show that glia are highly sensitive to partial loss of these genes as exemplified in Figure 5 for *PICALM*.

As endocytic vesicles are processed in the cell, the retromer is critical for protein recycling including cell surface receptors and rhodopsin (Wang et al., 2014). We observed reduced LD formation when retromer function was targeted via RNAi in both neurons and glia (Figure 3). However, neurodegeneration was observed when the retromer was lost in glia, but not neurons. This suggests a different role for the retromer in neurons and glia. In glia, the LRP1 receptor is critical for LD formation and the retromer is required for LRP1 recycling back to the membrane for efficient uptake of lipid particles (Stockinger et al., 2002). It also plays a critical role in Tau spreading (Rauch et al., 2020). Hence, loss of LD due to retromer dysfunction in glia would lead to LD loss and neurodegeneration, consistent with our observations. In mice CS6253 increased LRP1 in the hippocampus of APOE4 mice but it did not affect APOE3 mice which is not inconsistent with our data. Loss of retromer in neurons, may also lead to a progressive neurodegeneration but neurons may be less sensitive to this loss, as knockdown of *Vps26* in both glia and neurons causes more severe neuron loss than does knockdown of *Vps26* in glia alone (Wang et al., 2014; this study). Follow-up studies are needed to distinguish these hypotheses, but it is becoming increasingly evident that the retromer plays critical roles in the maintenance of neurons in AD (Muhammad et al., 2008).

Our model predicts that as ROS levels rise with age or other environmental factors, it becomes more difficult for glia to sequester peroxidated lipids, which promotes neurodegeneration. Thus, while approaches to induce uptake of lipids to remove ROS and amyloid from neurons or the extracellular space, it’s important to consider additional approaches to neutralize ROS early in disease to prevent glial death and eventual neurodegeneration. Our model also helps explain the non-linear relationship between amyloid burden and clinical severity of disease. Human Aβ expression induces neurodegeneration in *Drosophila* (Chouhan et al., 2016) and neurological and behavioral phenotypes in mice (Kobayashi and Chen, 2005; Oakley et al., 2006). Notably, production of low levels of ROS or Aβ alone causes neurotoxicity in a very slow and progressive manner. In flies, overexpression of Aβ42 causes neuronal death after approximately 45 days (Chouhan et al., 2016). However, we found that combining low amounts of ROS in Aβ42-expressing flies strongly exacerbated neurodegeneration and enhanced Aβ deposition in 5XFAD mice (Figure 6). It is noteworthy that Aβ and peroxidated lipids both bind APOE (Allan Butterfield et al., 2002; Lanfranco et al., 2020), providing a possible mechanism of ROS/Aβ42 synergy. Importantly, APOE4 is not properly lipidated and lipidation of APOE4 is required for Aβ42 binding (Kanekiyo et al., 2014; Namba et al., 1991; Strittmatter et al., 1993). Hence, APOE4 is unable to properly clear peroxidated lipids as well as Aβ42, strongly accelerating the demise of neurons. High amyloid burden may induce severe disease only in the presence of genetic or environmental triggers that induce ROS.

Regardless of the cause of ROS (e.g. age, environmental stress, or genetic perturbations), oxidative stress may initiate disease in an individual with a previous genetic predisposition to disease (APOE4, or other genetic disease risk). This model predicts that a reduction of oxidative stress, regardless of the cause, may help mitigate damage done by peroxidated lipids, and prevent further lipid biogenesis in neurons. Our model also suggest that the risk genes identified in GWAS that are involved in lipid handling and endocytosis may affect transport of peroxidated lipids from neurons into glia. Indeed, antioxidant levels are altered in AD patients and the use of antioxidants as a treatment for AD has been proposed previously, although with mixed outcomes (Frank and Gupta, 2005; Mullan et al., 2017; Vina et al., 2011; Wojtunik-Kulesza et al., 2016). Hence, animal models that better recapitulate AD phenotypes should consider the use of a combination of genetic manipulation and environmental ROS.

Numerous mammalian models have been generated to model AD that induce amyloid and/or tau production in the brain. Despite clear differences between the etiology and risk factors associated with the development of FAD and sporadic AD, many research groups model disease in animals using dominant FAD-causing mutations. This is because both forms of AD are histopathologically and clinically similar and because sporadic AD cannot be easily modelled in animals (LaFerla and Green, 2012). One historically common choice for mouse models of AD research is the 5XFAD mouse which harbors three mutations in the APP locus and two mutations in PS1 (Oakley et al., 2006). Each of these five mutations, on their own, are FAD-causing mutations in humans (Jankowsky and Zheng, 2017). Although this mouse line exhibits a very high amyloid burden, it almost certainly does not provide the most relevant model for human disease. While these mammalian models have proven fruitful for understanding many aspects of AD, none of them is adequate in reproducing the entirety of symptoms manifested in AD. We argue that the efficacy in disease modeling in mammals may be improved by the addition of ROS which is largely absent in the AD field currently. We acknowledge that the addition of ROS in mammalian models comes with various challenges including the use of toxic drugs (i.e. rotenone) or bulky and expensive equipment (i.e. hyperoxic animal chambers). Genetic mutations that induce ROS may be a more viable option to include in the background of AD models. A study using a mouse model of Leigh Syndrome in which the gene NDUFS4 is knocked out, thereby reducing activity of Complex I and leading to elevated ROS production results in early death of the Ndufs4^-/-^ animals (Assouline et al., 2012; Quintana et al., 2012). These mice have numerous LD in astrocytes and microglia prior to the onset of neuronal loss (Liu et al., 2015). In hypoxia these mice live much longer than when reared in normoxic conditions (Jain et al., 2016). Thus, the addition of ROS via genetic means, by for example removing a copy of *Ndufs4* may prove a viable method to induce ROS in an otherwise monogenic AD mammalian model.

Although age and mitochondrial dysfunction are obvious sources of ROS in AD patients, there may be numerous other conditions that induce ROS production and subsequent lipid peroxidation, LD formation, and eventual neurodegeneration. A careful examination of ROS in AD patients and inclusion of ROS in animal models may help begin to provide mechanistic insight into the etiology and progression of this complex disease. We argue that the use of antioxidants that penetrate the blood-brain barrier requires further investigation and that these treatments may aid prevention of neuron loss as observed in flies (Liu et al., 2015).

## Acknowledgements

We are grateful to the Bloomington Drosophila Stock Center and the Vienna Drosophila Resource Center for providing reagents. 5XFAD mice were provided to us through a gift of Huda Zoghbi. Mouse pathological studies were carried out by the Cell and Tissue Pathogenesis Core which is supported by the Eunice Kennedy Shriver National Institute of Child Health & Human Development of the National Institutes of Health (NIH) under the award number P50HD103555. Confocal images were captured in the Neurovisualization Core of the Intellectual and Developmental Disabilities Research Center (IDDRC), which is supported by the NIH under the award number U54HD083092. This work was supported by a grant from the Texas Alzheimer’s Research and Care Consortium under the award number 2018-05-11-II, to H.J.B. M.J.M. was supported by the Medical Genetics Research Fellowship Training Grant from the NIH under the award number T32 GM07526-41. L.D.G. was supported by the Brain Disorders & Development Fellowship Training Grant from the NIH under the award number T32 NS043124-18. P.C.M. is supported by a grant from Canadian Institutes of Health Research under the award number MFE-164712. M.S.I. is supported by the Canadian Institutes of Health Research under the award number 173321, and the Heart & Stroke Foundation of Canada under the award number 170722. I.R. is supported by the Alberta Synergies in Alzheimer’s and Related Disorders (SynAD) program funded by the Alzheimer Society of Alberta and Northwest Territories through their Hope for Tomorrow program and the University Hospital Foundation. SynAD operates in partnership with the Neuroscience and Mental Health Institute at the University of Alberta. H.J.B. is an investigator of the Howard Hughes Medical Institute (HHMI) and thanks HHMI for support.

## Author Contributions

Conceptualization, M.J.M., J.O.J., M.S.I. and H.J.B.; Investigation, M.J.M., S.B., J.G.H., P.C.M., I.R., J.C., and M.S.I.; Resources, J.O.J. and H.J.B.; Writing – Original Draft, M.J.M., J.G.H., and H.B.; Writing – Review & Editing, M.J.M., S.B., L.D.G., P.C.M., J.G.H., M.S.I, and H.J.B.; Supervision, H.J.B.; Funding Acquisition, H.J.B.

## Star Methods

### Resource Availability

#### Lead Contact

Further information and requests for resources and reagents should be directed to and will be fulfilled by the Lead Contact, Hugo Bellen.

#### Materials Availability

The peptide-expressing fly stock generated in this study was generated with permission through a collaboration with J.O.J and Artery Therapeutics, Inc. All stocks and reagents are freely available upon request to H.J.B.

#### Data and Code Availability

This study did not generate/analyze datasets.

### Experimental Model and Subject Details

#### Animals

*Drosophila melanogaster* were raised on standard molasses-based lab diet at 22°C under constant light conditions unless otherwise indicated. Stock genotypes and availability is listed in the Key Resources Table. Female flies were used for all experiments.

Experiments using mice, *Mus musculus*, were carried out under the approval of the Animal Care and Use Committee at Baylor College of Medicine. Mice were housed under 12-hour light/dark conditions at 22°C with standard chow diet (5053; Picolab) and water available ad libitum. Stock genotypes and availability is listed in the Key Resources Table. Hyperoxia experiments were carried using the A-Chamber animal cage enclosure (BioSpherix) with oxygen levels regulated using the ProOx360 High Infusion Rate O2 Controller (BioSpherix).

### Method Details

#### Gene Tree Assembly

Protein sequences of all human and fly ABCA genes were downloaded from the National Center for Biotechnology Information (NCBI) website. Sequences were aligned using ClustalW algorithm within MEGA X (Kumar et al., 2018). Aligned sequences were used to build the neighbor-joining tree within MEGA X.

#### Generation of Transgenic Flies

UAS-ArgosSS::Peptide transgenic flies were generated by ORF synthesis in pUC57 using Drosophila codon-optimized sequence (Integrated DNA Technologies) of the argos secretion signal (MPTTLMLLPCMLLLLLTAAAVAVGG) (Chouhan et al., 2016) upstream of the peptide sequence, where citrulline residues were replaced with arginine residues, (EVRSKLEEWLAALRELAEELLARAKS) (Bielicki, 2016; Boehm-Cagan and Michaelson, 2014). The ORF was shuttled to the Gateway pDONR221 entry vector (ThermoFisher) by BP clonase II reaction (ThermoFisher) using Argos_attB primers. Fully sequence verified clones were shuttled to the pGW-attB-HA destination vector (Bischof et al., 2012) by LR clonase II (ThermoFisher). The UAS construct was inserted into the VK37 (PBac{y[+]-attP}VK00037) docking site by ϕC31 mediated transgenesis (Venken et al., 2006).

#### Lipid Droplet Analysis

Whole-mount staining of fly retinas with Nile Red to visualize lipids was performed as in Liu, et al. (2015). In brief, fly heads were isolated under PBS and fixed in 3.7% formaldehyde overnight. Retinas were then dissected under PBS and rinsed three times with 1X PBS and incubated for 20 minutes at 1:1,000 dilution of PBS with 1 mg/mL Nile Red (Millipore Sigma). Retinas were subsequently rinsed five times with 1X PBS and mounted in Vectashield (Vector Labs) for imaging on a Leica SP8 confocal microscope. Images were obtained using a 63X glycerol submersion lens with 3X zoom.

#### Electroretinogram Assay

Electroretinogram (ERG) assays (Heisenberg, 1971) were performed as previously described (Verstreken et al., 2003). In brief, live flies were immobilized with Elmer’s school glue on a microscope slide. Glass electrodes, filled with 3 M NaCl, were placed in the thorax for reference and on the center part of the eye for recording. Prior to recording, flies were maintained in the dark for at least one minute. Approximately one-second light flashes were manually delivered using a halogen lamp. At least three recordings from at least 10 flies per genotype were obtained for analysis using LabChart 8.

#### Immunohistochemistry

Animal perfusion, sectioning and immunohistochemistry was performed as in Sillitoe, et al. (2008). In brief, mice were anesthetized with and sacrificed by intracardiac perfusion, initially with saline, followed by 4% paraformaldehyde in PBS. Mice brains were removed and bisected down the midline with one hemibrain utilized for histopathological analysis, which was immersed again in paraformaldehyde for 24 hours at 4°C. Hemibrains were then dehydrated and preserved in paraffin. Serial sagittal sections were cut and 6 µM sections were mounted on glass slides. After sections were blocked in 10% normal goat serum and rinsed with PBS, sections were stained with the anti-Aβ42 antibody (Covance, 1:1,000) and a biotinylated anti-mouse IgG antibody (Jackson, 1:200). Slides were then incubated in DAB solution, monitored by eye, and the reaction stopped with distilled water. Finally, slides were counterstained using Mayer’s hematoxylin and dehydrated prior to imaging.

#### qRT-PCR Analysis

RNA extraction, cDNA synthesis, and qRT-PCR were performed as in Barish et al. (2018). In brief, 10 larvae ubiquitously expressing RNAi (via Daugtherless-Gal4) were isolated. RNA extraction was carried out using the RNeasy Mini Kit (Qiagen) followed by reverse transcription into cDNA using iScript Reverse Transcription Supermix (BioRad). Quantitative PCR reactions were performed using iTaq Universal SYBR Green Supermix (BioRad) on a BioRad CFX96 Touch Real-Time PCR Detection System. Three biological and technical replicates were performed for each genotype. Expression (Ct) values were obtained and used to calculate differential expression (ΔCt) and normalized to GAPDH expression.

#### Primary culture of hippocampal neurons and astrocytes

Hippocampal cultures were generated from P0-P1 Sprague-Dawley rats obtained from Charles River Laboratories that arrived at our facility one week prior to birth. These experiments were approved by the Canadian Council of Animal Care at the University of Alberta (AUP#3358). Cultures were prepared as previously described (Beaudoin et al., 2012; Ioannou et al., 2019a). In brief, tissue was digested with papain, gently triturated, and filtered with a cell strainer and plated on poly-D-lysine coated coverslips for the transfer assay or plastic tissue culture dishes for Western blot analysis. Neurons were grown in Neurobasal medium containing B-27 supplement, 2 mM Glutamax and antibiotic-antimycotic. Astrocytes were grown in Basal Eagle Media containing 10% fetal bovine serum, 0.45% glycose, 1 mM sodium pyruvate, 2 mM Glutamax, and antibiotic-antimycotic. All cells were grown at 37°C in 5% CO_2_.

#### Lentivirus transduction

Astrocytes at DIV 2 were transduced with SMARTVector lentiviral shRNA (Dharmacon) at an MOI of 3. Three independent shRNA sequences targeting PICALM or a non-targeting control shRNA was used. The media was replaced with fresh culture media after 24 hrs. and the cells were used for protein validation or in the transfer assay 5 days later (DIV 7). To validate protein knockdown, astrocytes were lysed in lysis buffer [20 mM HEPES pH 7.4, 100 mM NaCl, 1% Triton X-100, 5 mM EDTA, 1× Halt Protease & Phosphatase Inhibitor Cocktail (Thermo Scientific)], resolved by SDS–PAGE and processed for Western blotting using anti-PICALM rabbit polyclonal (Millipore Sigma) and anti-GAPDH mouse monoclonal (ThermoFisher) as a loading control. Lysates were run in duplicate and statistics were performed on the average of these duplicates for each experiment.

#### Fatty acid transfer assay

Neurons (DIV 7) were incubated with 2 μM BODIPY 558/568 (Red-C12) for 16 hours in neuronal growth media. Neurons were washed twice with warm phosphate-buffered saline (PBS) and incubated with fresh media for 1 hour. Red-C12 labelled neurons and unlabeled astrocytes transduced with lentivirus as described above were washed twice with warm PBS and the coverslips were cultured together (facing each other), separated by paraffin wax and incubated in Hanks’ Balanced Salt solution containing calcium and magnesium for 4 hours at 37°C (Ioannou et al., 2019b, 2019a). Astrocytes were fixed in 4% paraformaldehyde, stained with DAPI, and mounted using DAKO fluorescence mounting media. Images were acquired using a Zeiss 710 Laser Scanning Confocal Microscope equipped with a plan-apochromat 63x oil objective (Zeiss, NA = 1.4). Maximum intensity projections of three-dimensional image stacks (0.5µm sections) of Red-C12 staining were obtained and analyzed using ImageJ. Only astrocytes expressing the lentiviral reporter turboGFP were quantified. Images were thresholded and the number of particles with a pixel size greater than 2 was detected. 10 cells per coverslip were averaged. Schematic of fatty acid transfer assay was created with BioRender.com.

### Quantification and Statistical Analysis

FIJI (Schindelin et al., 2012) was utilized to view fly retinal and mouse brain images and all genotypes were blinded prior to quantification. Lipid droplets with diameter ≥0.5 µM were manually quantified from fly retinal images. Amyloid plaque number from mouse brain images was manually quantified and amyloid size measurements were taken using the ‘Measure’ tool in FIJI. LabChart 8 (AD Instruments) was used to view and measure the amplitude of ERG traces. Quantification datasets were assembled in Microsoft Excel 365 for comparison and statistical analysis. For quantification, ≥10 animals per genotype were used. Mean +/− SEM were plotted and pair-wise T-tests were performed with a statistical significance cutoff at *p<0.05, and **p<0.01. Statistical analysis of knockdown efficiency in rat cells used the Kruskal-Wallis test with Dunn’s posttest using a significance cutoff at *p<0.05. Analysis of lipid transfer utilized One-way ANOVA with Dunnett’s posttest using a significance cutoff at *** p < 0.001.

### KEY RESOURCES TABLE

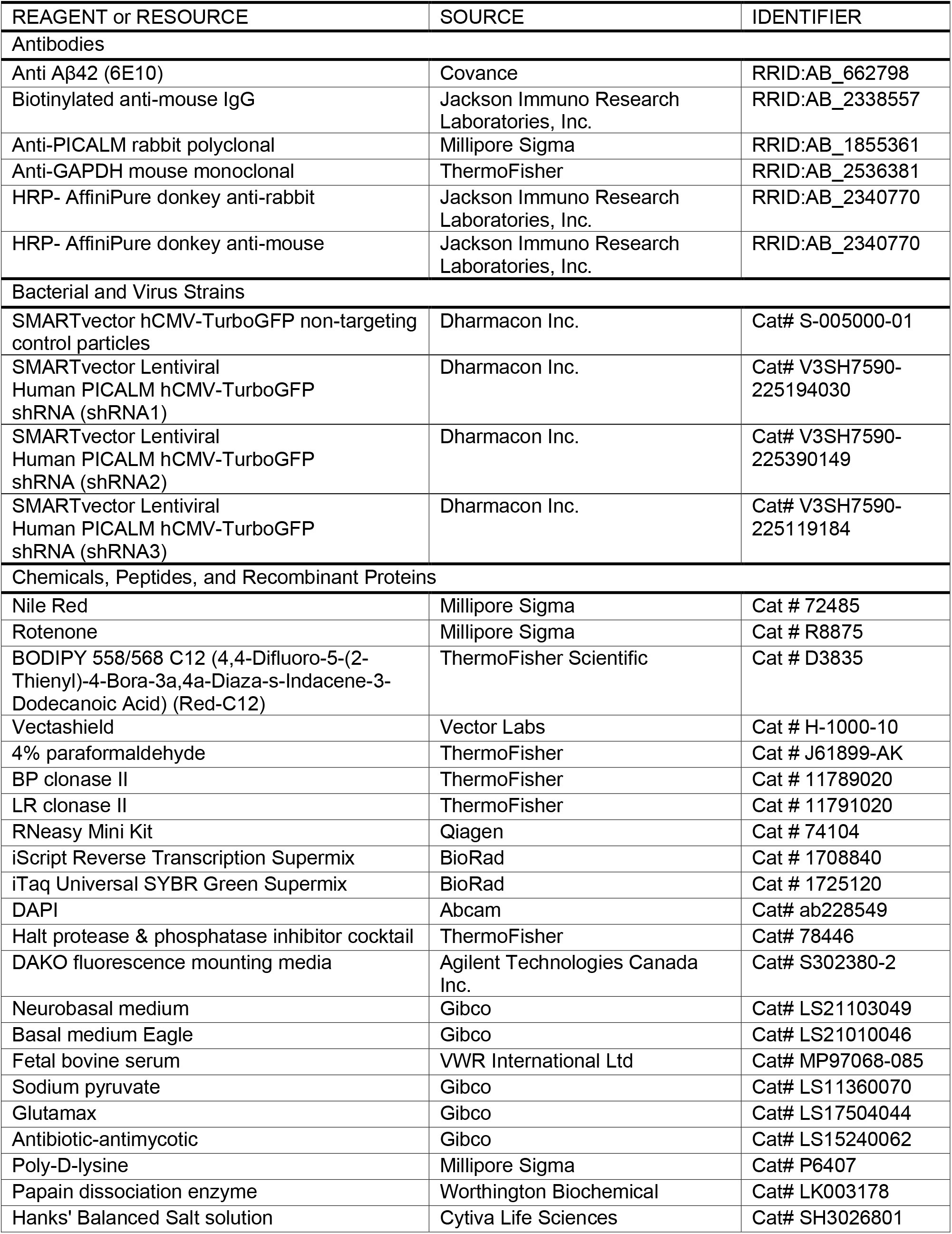
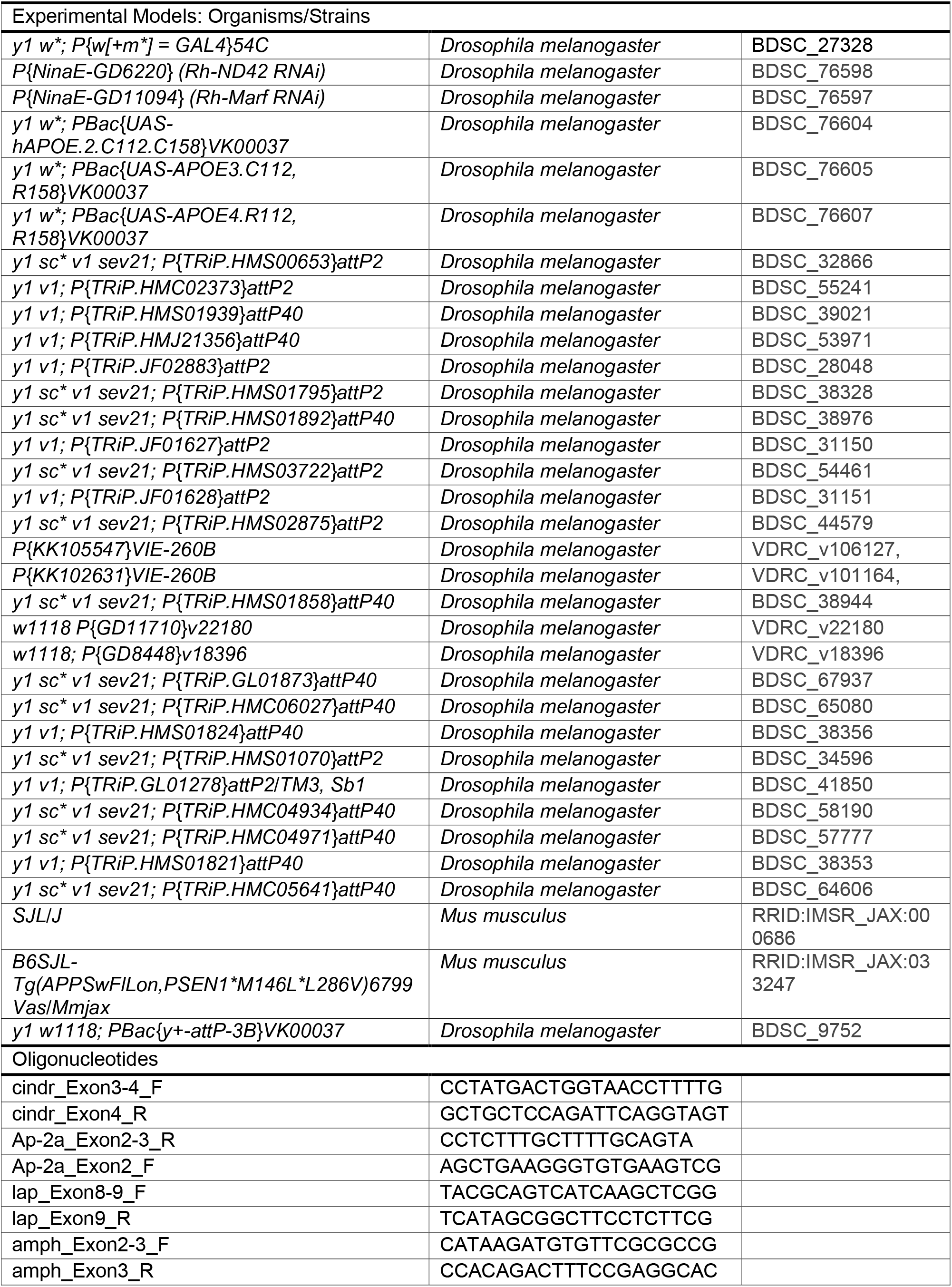
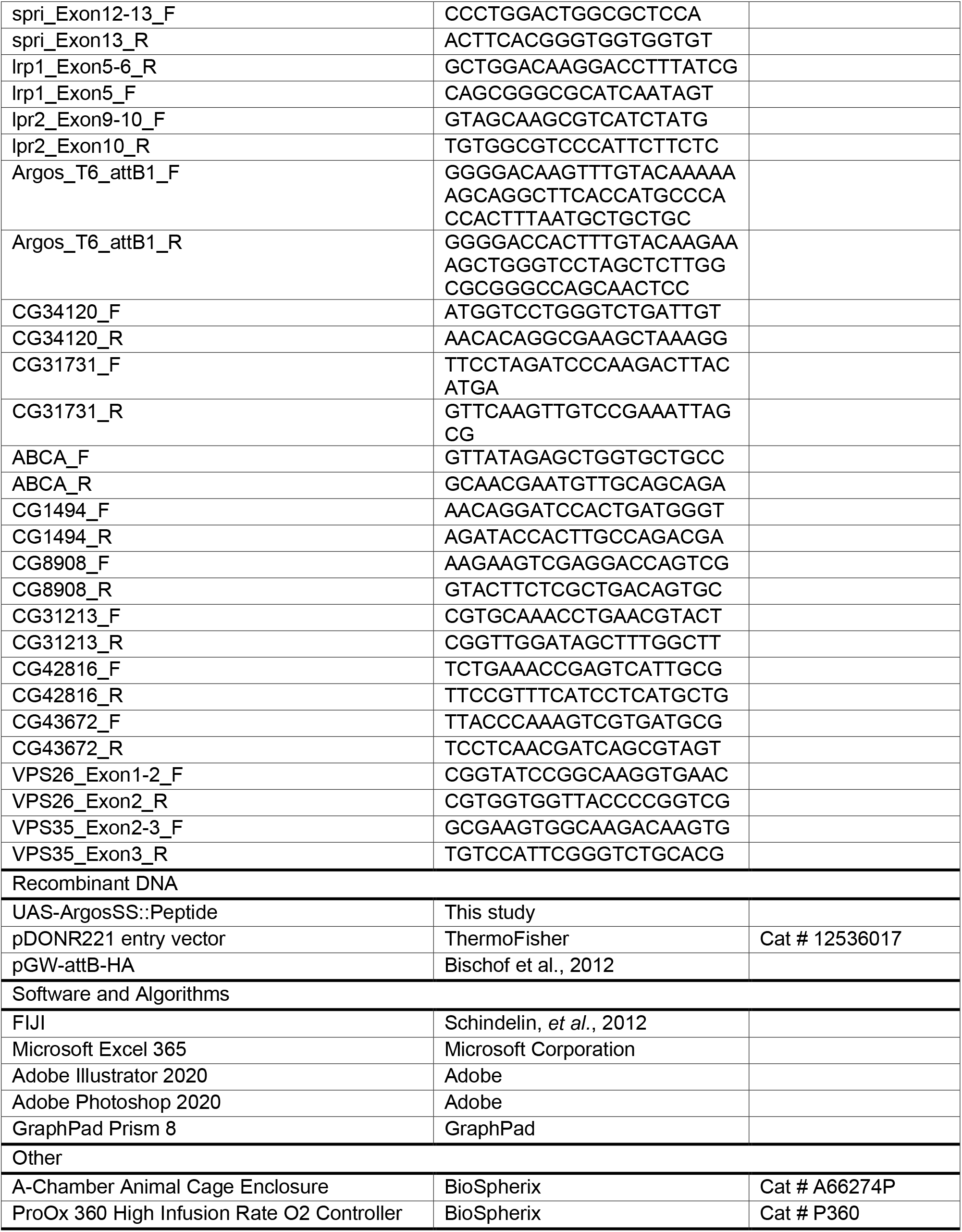

---

**Supplemental Figure 1.**
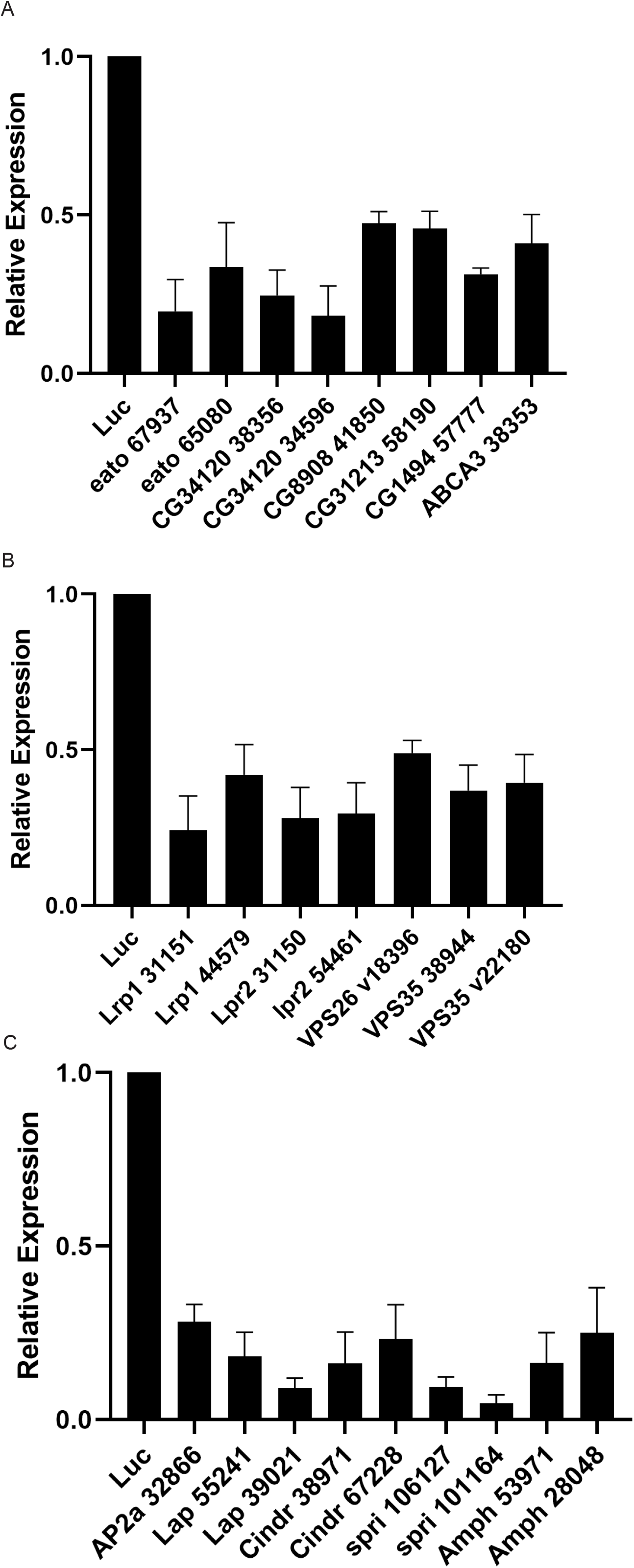
Analysis of RNA expression of genes targeted in this study. (A-C) Quantitation of mRNA expression of the genes assayed after RNAi induction in fly heads. Animals are housed at 29°C under 12-hour light/dark conditions. In each assay, a luciferase RNAi is used as a negative control. RNAi induces at least a 50% reduction of the mRNA of all genes assayed.

## References

A. Alzheimer (1911). On certain peculiar diseases of old age. Clin. Anat. 8, 429–431.

Abe-Dohmae, S., Ikeda, Y., Matsuo, M., Hayashi, M., Okuhira, K.I., Ueda, K., and Yokoyama, S. (2004). Human ABCA7 Supports Apolipoprotein-mediated Release of Cellular Cholesterol and Phospholipid to Generate High Density Lipoprotein. J. Biol. Chem. 279, 604–611.

Abuznait, A.H., and Kaddoumi, A. (2012). Role of ABC transporters in the pathogenesis of Alzheimers disease. ACS Chem. Neurosci. 3, 820–831.

Van Acker, Z.P., Bretou, M., and Annaert, W. (2019). Endo-lysosomal dysregulations and late-onset Alzheimer’s disease: Impact of genetic risk factors. Mol. Neurodegener. 14, 1–20.

Aikawa, T., Holm, M.L., and Kanekiyo, T. (2018). ABCA7 and pathogenic pathways of Alzheimer’s disease. Brain Sci. 8, 1–13.

Allan Butterfield, D., and Boyd-Kimball, D. (2018). Oxidative Stress, Amyloid-β Peptide, and Altered Key Molecular Pathways in the Pathogenesis and Progression of Alzheimer’s Disease. J. Alzheimer’s Dis. 62, 1345–1367.

Allan Butterfield, D., Castegna, A., Lauderback, C.M., and Drake, J. (2002). Evidence that amyloid beta-peptide-induced lipid peroxidation and its sequelae in Alzheimer’s disease brain contribute to neuronal death. Neurobiol. Aging 23, 655–664.

Alzheimer, A. (1907). Uber eine eigenartige Erkrankung der Hirnrinde. Allg. Zeitschrift Rsychiatrie Psych. Medizine 64, 146–148.

Andreone, B.J., Przybyla, L., Llapashtica, C., Rana, A., Davis, S.S., van Lengerich, B., Lin, K., Shi, J., Mei, Y., Astarita, G., et al. (2020). Alzheimer’s-associated PLCγ2 is a signaling node required for both TREM2 function and the inflammatory response in human microglia. Nat. Neurosci. 23, 927–938.

Assouline, Z., Jambou, M., Rio, M., Bole-Feysot, C., de Lonlay, P., Barnerias, C., Desguerre, I., Bonnemains, C., Guillermet, C., Steffann, J., et al. (2012). A constant and similar assembly defect of mitochondrial respiratory chain complex I allows rapid identification of NDUFS4 mutations in patients with Leigh syndrome. 1822, 1062–1069.

Baig, S., Joseph, S.A., Tayler, H., Abraham, R., Owen, M.J., Williams, J., Kehoe, P.G., and Love, S. (2010). Distribution and expression of picalm in alzheimer disease. J. Neuropathol. Exp. Neurol. 69, 1071–1077.

Barish, S., Nuss, S., Strunilin, I., Bao, S., Mukherjee, S., Jones, C.D., and Volkan, P.C. (2018). Combinations of DIPs and Dprs control organization of olfactory receptor neuron terminals in Drosophila. PLoS Genet. 14, 1–33.

Beaudoin, G.M.J., Lee, S.H., Singh, D., Yuan, Y., Ng, Y.G., Reichardt, L.F., and Arikkath, J. (2012). Culturing pyramidal neurons from the early postnatal mouse hippocampus and cortex. Nat. Protoc. 7, 1741–1754.

Behl, C. (1997). Amyloid β-protein toxicity and oxidative stress in Alzheimer’s disease. Cell Tissue Res. 290, 471–480.

Berbée, J.F.P., Mol, I.M., Milne, G.L., Pollock, E., Hoeke, G., Lütjohann, D., Monaco, C., Rensen, P.C.N., van der Ploeg, L.H.T., and Shchepinov, M.S. (2017). Deuterium-reinforced polyunsaturated fatty acids protect against atherosclerosis by lowering lipid peroxidation and hypercholesterolemia. Atherosclerosis 264, 100–107.

Berman, D.E., Ringe, D., Petsko, G.A., and Small, S.A. (2015). The Use of Pharmacological Retromer Chaperones in Alzheimer’s Disease and other Endosomal-related Disorders. Neurotherapeutics 12, 12–18.

Bielicki, J.K. (2016). ABCA1 agonist peptides for the treatment of disease. Curr. Opin. Lipidol. 27, 40–46.

Bischof, J., Björklund, M., Furger, E., Schertel, C., Taipale, J., and Basler, K. (2012). A versatile platform for creating a comprehensive UAS-ORFeome library in Drosophila. Dev. 140, 2434–2442.

Bisht, K., Sharma, K., and Tremblay, M.È. (2018). Chronic stress as a risk factor for Alzheimer’s disease: Roles of microglia-mediated synaptic remodeling, inflammation, and oxidative stress. Neurobiol. Stress 9, 9–21.

Bloom, G.S. (2014). Amyloid-β and tau: The trigger and bullet in Alzheimer disease pathogenesis. JAMA Neurol. 71, 505–508.

Boehm-Cagan, A., and Michaelson, D.M. (2014). Reversal of apoE4-Driven Brain Pathology and Behavioral Deficits by Bexarotene. J. Neurosci. 34, 7293–7301.

Boehm-Cagan, A., Bar, R., Liraz, O., Bielicki, J.K., Johansson, J.O., and Michaelson, D.M. (2016a). ABCA1 Agonist Reverses the ApoE4-Driven Cognitive and Brain Pathologies. J. Alzheimer’s Dis. 54, 1219–1233.

Boehm-Cagan, A., Bar, R., Harats, D., Shaish, A., Levkovitz, H., Bielicki, J.K., Johansson, J.O., and Michaelson, D.M. (2016b). Differential Effects of apoE4 and Activation of ABCA1 on Brain and Plasma Lipoproteins. PLoS One 11, 1–17.

Bossche, T. Van Den, Sleegers, K., Cuyvers, E., Engelborghs, S., Sieben, A., Roeck, A. De Cauwenberghe, C. Van, Vermeulen, S., Broeck, M. Van Den, Laureys, A., et al. (2016). Phenotypic characteristics of Alzheimer patients carrying an ABCA7 mutation. Neurology 86, 2126–2133.

Bradley-Whitman, M.A., and Lovell, M.A. (2015). Biomarkers of lipid peroxidation in Alzheimer disease (AD): an update. Arch. Toxicol. 89, 1035–1044.

Bradley, M.A., Xiong-Fister, S., Markesbery, W.R., and Lovell, M.A. (2012). Elevated 4-hydroxyhexenal in Alzheimer’s disease (AD) progression. Neurobiol. Aging 33, 1034–1044.

Van Den Brink, D.M., Cubizolle, A., Chatelain, G., Davoust, N., Girard, V., Johansen, S., Napoletano, F., Dourlen, P., Guillou, L., Angebault-Prouteau, C., et al. (2018). Physiological and pathological roles of FATP-mediated lipid droplets in Drosophila and mice retina. PLOS Genet. 14, 1–25.

Burnside, S.W., and Hardingham, G.E. (2017). Transcriptional regulators of redox balance and other homeostatic processes with the potential to alter neurodegenerative disease trajectory.

Butterfield, D.A. (2020). Brain lipid peroxidation and alzheimer disease: Synergy between the Butterfield and Mattson laboratories. Ageing Res. Rev. 64, 1568–1637.

Chang, Y.T., Hsu, S.W., Huang, S.H., Huang, C.W., Chang, W.N., Lien, C.Y., Lee, J.J., Lee, C.C., and Chang, C.C. (2019). ABCA7 polymorphisms correlate with memory impairment and default mode network in patients with APOEɛ4-associated Alzheimer’s disease. Alzheimer’s Res. Ther. 11, 103–113.

Chen, Q., Liang, B., Wang, Z., Cheng, X., Huang, Y., Liu, Y., and Huang, Z. (2016). Influence of four polymorphisms in ABCA1 and PTGS2 genes on risk of Alzheimer’s disease: a meta-analysis. Neurol. Sci. 37, 1209–1220.

Chouhan, A.K., Guo, C., Hsieh, Y.-C.C., Ye, H., Senturk, M., Zuo, Z., Li, Y., Chatterjee, S., Botas, J., Jackson, G.R., et al. (2016). Uncoupling neuronal death and dysfunction in Drosophila models of neurodegenerative disease. Acta Neuropathol. Commun. 4, 62–76.

Chung, H. lok, Wangler, M.F., Marcogliese, P.C., Jo, J., Ravenscroft, T.A., Zuo, Z., Duraine, L., Sadeghzadeh, S., Li-Kroeger, D., Schmidt, R.E., et al. (2020). Loss- or Gain-of-Function Mutations in ACOX1 Cause Axonal Loss via Different Mechanisms. Neuron 106, 589–606.e6.

Conejero-Goldberg, C., Gomar, J.J., Bobes-Bascaran, T., Hyde, T.M., Kleinman, J.E., Herman, M.M., Chen, S., Davies, P., and Goldberg, T.E. (2014). APOE2 enhances neuroprotection against alzheimer’s disease through multiple molecular mechanisms. Mol. Psychiatry 19, 1243–1250.

Dhawan, K., Naslavsky, N., Caplan, S., and Hanson, P.I. (2020). Sorting nexin 17 (SNX17) links endosomal sorting to Eps15 homology domain protein 1 (EHD1)-mediated fission machinery. J. Biol. Chem. 295, 3837–3850.

Ermondi, G., Catalano, F., Vallaro, M., Ermondi, I., Leal, M.P.C., Rinaldi, L., Visentin, S., and Caron, G. (2015). Lipophilicity of amyloid β-peptide 12-28 and 25-35 to unravel their ability to promote hydrophobic and electrostatic interactions. Int. J. Pharm. 495, 179–185.

Fan, J., Donkin, J., and Wellington, C. (2009). Greasing the wheels of Aβ clearance in Alzheimer’s Disease: The role of lipids and apolipoprotein e. BioFactors 35, 239–248.

Fehér, Á., Giricz, Z., Juhász, A., Pákáski, M., Janka, Z., and Kálmán, J. (2018). ABCA1 rs2230805 and rs2230806 common gene variants are associated with Alzheimer’s disease. Neurosci. Lett. 664, 79–83.

Ferrari, M., Jain, I.H., Goldberger, O., Rezoagli, E., Thoonen, R., Cheng, K.-H., Sosnovik, D.E., Scherrer-Crosbie, M., Mootha, V.K., and Zapol, W.M. (2017). Hypoxia treatment reverses neurodegenerative disease in a mouse model of Leigh syndrome. Proc. Natl. Acad. Sci. U. S. A. 114, E4241–E4250.

Frank, B., and Gupta, S. (2005). A review of antioxidants and Alzheimer’s disease. Ann. Clin. Psychiatry 17, 269–286.

Furusawa, K., Takasugi, T., Chiu, Y.W., Hori, Y., Tomita, T., Fukuda, M., and Hisanaga, S. ichi (2019). CD2-associated protein (CD2AP) overexpression accelerates amyloid precursor protein (APP) transfer from early endosomes to the lysosomal degradation pathway. J. Biol. Chem. 294, 10886–10899.

Gan, Q., and Watanabe, S. (2018). Synaptic vesicle endocytosis in different model systems. Front. Cell. Neurosci. 12, 171–197.

Giau, V. Van, Bagyinszky, E., Yang, Y.S., Youn, Y.C., An, S.S.A., and Kim, S.Y. (2019). Genetic analyses of early-onset Alzheimer’s disease using next generation sequencing. Sci. Rep. 9, 1–10.

González-Gaitán, M., and Jäckle, H. (1997). Role of drosophila α-adaptin in presynaptic vesicle recycling. Cell 88, 767–776.

Götz, J., Deters, N., Doldissen, A., Bokhari, L., Ke, Y., Wiesner, A., Schonrock, N., and Ittner, L.M. (2007). A decade of tau transgenic animal models and beyond. In Brain Pathology, (John Wiley & Sons, Ltd), pp. 91–103.

Götz, J., Eckert, A., Matamales, M., Ittner, L.M., and Liu, X. (2011). Modes of Aβ toxicity in Alzheimer’s disease. Cell. Mol. Life Sci. 68, 3359–3375.

Griendling, K.K., Touyz, R.M., Zweier, J.L., Dikalov, S., Chilian, W., Chen, Y.R., Harrison, D.G., and Bhatnagar, A. (2016). Measurement of Reactive Oxygen Species, Reactive Nitrogen Species, and Redox-Dependent Signaling in the Cardiovascular System: A Scientific Statement from the American Heart Association. Circ. Res. 119, e39–e75.

Grimm, A., and Eckert, A. (2017). Brain aging and neurodegeneration: from a mitochondrial point of view. J. Neurochem. 143, 418–431.

Hafiane, A., Bielicki, J.K., Johansson, J.O., and Genest, J. (2015). Novel apo E-derived ABCA1 agonist peptide (CS-6253) promotes reverse cholesterol transport and induces formation of preβ-1 HDL in vitro. PLoS One 10, 1–32.

Hardy, J., Selkoe, D.J., Ovod, V., Munsell, L., Kasten, T., Morris, J.C., Yarasheski, K.E., and Bateman, R.J. (2002). The amyloid hypothesis of Alzheimer’s disease: progress and problems on the road to therapeutics. Science 297, 353–356.

Harrison, B.J., Venkat, G., Lamb, J.L., Hutson, T.H., Drury, C., Rau, K.K., Bunge, M.B., Mendell, L.M., Gage, F.H., Johnson, R.D., et al. (2016). The adaptor protein CD2AP is a coordinator of neurotrophin signaling-mediated axon arbor plasticity. J. Neurosci. 36, 4259–4275.

Hatters, D.M., Peters-Libeu, C.A., and Weisgraber, K.H. (2006). Apolipoprotein E structure: insights into function. Trends Biochem. Sci. 31, 445–454.

Hayashi, M., Abe-Dohmae, S., Okazaki, M., Ueda, K., and Yokoyama, S. (2005). Heterogeneity of high density lipoprotein generated by ABCA1 and ABCA7. J. Lipid Res. 46, 1703–1711.

Heisenberg, M. (1971). Separation of Receptor and Lamina Potentials in the Electroretinogram of Normal and Mutant Drosophila. J. Exp. Biol. 55, 85LP–100.

Herz, J. (2009). Apolipoprotein E receptors in the nervous system. Curr. Opin. Lipidol. 20, 190–196.

Huang, Y., and Mahley, R.W. (2014). Apolipoprotein E: Structure and function in lipid metabolism, neurobiology, and Alzheimer’s diseases. Neurobiol. Dis. 72, 3–12.

Huynh, T.-P.V.P. V., Davis, A.A., Ulrich, J.D., and Holtzman, D.M. (2017). Apolipoprotein E and Alzheimer’s disease: the influence of apolipoprotein E on amyloid-? and other amyloidogenic proteins. 58, 824–836.

Ioannou, M.S., Jackson, J., Sheu, S.-H., Chang, C.-L., Weigel, A. V., Liu, H., Pasolli, H.A., Xu, C.S., Pang, S., Matthies, D., et al. (2019a). Neuron-Astrocyte Metabolic Coupling Protects against Activity-Induced Fatty Acid Toxicity. Cell 177, 1522–1535.e14.

Ioannou, M.S., Liu, Z., and Lippincott-Schwartz, J. (2019b). A Neuron-Glia Co-culture System for Studying Intercellular Lipid Transport. Curr. Protoc. Cell Biol. 84, 1–21.

Jain, I.H., Zazzeron, L., Goli, R., Alexa, K., Schatzman-Bone, S., Dhillon, H., Goldberger, O., Peng, J., Shalem, O., Sanjana, N.E., et al. (2016). Hypoxia as a therapy for mitochondrial disease. Science (80-.). 352, 54–61.

Jaiswal, M., Sandoval, H., Zhang, K., Bayat, V., and Bellen, H.J. (2012). Probing mechanisms that underlie human neurodegenerative diseases in Drosophila. Annu. Rev. Genet. 46, 371–396.

Jankowsky, J.L., and Zheng, H. (2017). Practical considerations for choosing a mouse model of Alzheimer’s disease. Mol. Neurodegener. 12, 89–110.

Kaksonen, M., and Roux, A. (2018). Mechanisms of clathrin-mediated endocytosis. Nat. Rev. Mol. Cell Biol. 19, 313–326.

Kanekiyo, T., Xu, H., and Bu, G. (2014). ApoE and Aβ in Alzheimer’s disease: Accidental encounters or partners? Neuron 81, 740–754.

Karch, C.M., and Goate, A.M. (2015). Alzheimer’s disease risk genes and mechanisms of disease pathogenesis. Biol. Psychiatry 77, 43–51.

Kobayashi, D.T., and Chen, K.S. (2005). Behavioral phenotypes of amyloid-based genetically modified mouse models of Alzheimer’s disease. Genes, Brain Behav. 4, 173–196.

Koldamova, R., Staufenbiel, M., and Lefterov, I. (2005). Lack of ABCA1 considerably decreases brain ApoE level and increases amyloid deposition in APP23 mice. J. Biol. Chem. 280, 43224–43235.

Kumar, S., Stecher, G., Li, M., Knyaz, C., and Tamura, K. (2018). MEGA X: Molecular evolutionary genetics analysis across computing platforms. Mol. Biol. Evol. 35, 1547–1549.

Kunkle, B.W., Grenier-Boley, B., Sims, R., Bis, J.C., Damotte, V., Naj, A.C., Boland, A., Vronskaya, M., van der Lee, S.J., Amlie-Wolf, A., et al. (2019). Genetic meta-analysis of diagnosed Alzheimer’s disease identifies new risk loci and implicates Aβ, tau, immunity and lipid processing. Nat. Genet. 51, 414–430.

LaFerla, F.M., and Green, K.N. (2012). Animal models of Alzheimer disease. Cold Spring Harb. Perspect. Med. 2, 1–13.

Lambert, J.-C., Ibrahim-Verbaas, C.A., Harold, D., Naj, A.C., Sims, R., Bellenguez, C., Jun, G., DeStefano, A.L., Bis, J.C., Beecham, G.W., et al. (2013). Meta-analysis of 74,046 individuals identifies 11 new susceptibility loci for Alzheimer’s disease. Nat. Genet. 45, 1452–1458.

Lane-Donovan, C., and Herz, J. (2017). ApoE, ApoE Receptors, and the Synapse in Alzheimer’s Disease. Trends Endocrinol. Metab. 28, 273–284.

Lanfranco, M.F., Ng, C.A., and Rebeck, G.W. (2020). ApoE lipidation as a therapeutic target in Alzheimer’s disease. Int. J. Mol. Sci. 21, 1–19.

Lee, E., Marcucci, M., Daniell, L., Pypaert, M., Weisz, O.A., Ochoa, G.C., Farsad, K., Wenk, M.R., and De Camilli, P. (2002). Amphiphysin 2 (Bin1) and T-tubule biogenesis in muscle. Science (80-.). 297, 1193–1196.

Lee, P.T., Zirin, J., Kanca, O., Lin, W.W., Schulze, K.L., Li-Kroeger, D., Tao, R., Devereaux, C., Hu, Y., Chung, V., et al. (2018). A gene-specific T2A-GAL4 library for drosophila. Elife 7.

Li, Z., Shue, F., Zhao, N., Shinohara, M., and Bu, G. (2020). APOE2: protective mechanism and therapeutic implications for Alzheimer’s disease. Mol. Neurodegener. 15, 63–81.

Lin, G., Wang, L., Marcogliese, P.C., and Bellen, H.J. (2019). Sphingolipids in the Pathogenesis of Parkinson’s Disease and Parkinsonism. Trends Endocrinol. Metab. 30, 106–117.

Liu, L., Zhang, K., Sandoval, H., Yamamoto, S., Jaiswal, M., Sanz, E., Li, Z., Hui, J., Graham, B.H., Quintana, A., et al. (2015). Glial Lipid Droplets and ROS Induced by Mitochondrial Defects Promote Neurodegeneration. Cell 160, 177–190.

Liu, L., MacKenzie, K.R., Putluri, N., Maletić-Savatić, M., and Bellen, H.J. (2017). The Glia-Neuron Lactate Shuttle and Elevated ROS Promote Lipid Synthesis in Neurons and Lipid Droplet Accumulation in Glia via APOE/D. Cell Metab. 26, 719–737.e6.

Marioni, R.E., Harris, S.E., Zhang, Q., McRae, A.F., Hagenaars, S.P., Hill, W.D., Davies, G., Ritchie, C.W., Gale, C.R., Starr, J.M., et al. (2018). GWAS on family history of Alzheimer’s disease. Transl. Psychiatry 8, 1–7.

Masdeu, J.C. (2020). Neuroimaging of Diseases Causing Dementia. Neurol. Clin. 38, 65–94.

Michaelson, D.M. (2014). *APOE* ε4: The most prevalent yet understudied risk factor for Alzheimer’s disease. Alzheimer’s Dement. 10, 861–868.

Mietelska-Porowska, A., Wasik, U., Goras, M., Filipek, A., and Niewiadomska, G. (2014). Tau Protein Modifications and Interactions: Their Role in Function and Dysfunction. Int. J. Mol. Sci. 15, 4671–4713.

Moreira, P.I., Nunomura, A., Honda, K., Aliev, G., Casadesus, G., Zhu, X., Smith, M.A., and Perry, G. (2007). The key role of oxidative stress in alzheimer’s disease. In Oxidative Stress and Neurodegenerative Disorders, (Elsevier), pp. 267–281.

Muhammad, A., Flores, I., Zhang, H., Yu, R., Staniszewski, A., Planel, E., Herman, M., Ho, L., Kreber, R., Honig, L.S., et al. (2008). Retromer deficiency observed in Alzheimer’s disease causes hippocampal dysfunction, neurodegeneration, and Aβ accumulation. Proc. Natl. Acad. Sci. U. S. A. 105, 7327–7332.

Mullan, K., Williams, M.A., Cardwell, C.R., McGuinness, B., Passmore, P., Silvestri, G., Woodside, J. V., and McKay, G.J. (2017). Serum concentrations of vitamin E and carotenoids are altered in Alzheimer’s disease: A case-control study. Alzheimer’s Dement. Transl. Res. Clin. Interv. 3, 432–439.

Namba, Y., Tomonaga, M., Kawasaki, H., Otomo, E., and Ikeda, K. (1991). Apolipoprotein E immunoreactivity in cerebral amyloid deposits and neurofibrillary tangles in Alzheimer’s disease and kuru plaque amyloid in Creutzfeldt-Jakob disease. Brain Res. 541, 163–166.

Nelson, P.T., Fardo, D.W., and Katsumata, Y. (2020). The MUC6/AP2A2 Locus and Its Relevance to Alzheimer’s Disease: A Review. J. Neuropathol. Exp. Neurol. 79, 568–584.

Neumann, J., Rose-Sperling, D., and Hellmich, U.A. (2017). Diverse relations between ABC transporters and lipids: An overview. Biochim. Biophys. Acta - Biomembr. 1859, 605–618.

Nordestgaard, L.T., Tybjærg-Hansen, A., Nordestgaard, B.G., and Frikke-Schmidt, R. (2015). Loss-of-function mutation in ABCA1 and risk of Alzheimer’s disease and cerebrovascular disease. Alzheimer’s Dement. 11, 1430–1438.

Nugent, A.A., Lin, K., van Lengerich, B., Lianoglou, S., Przybyla, L., Davis, S.S., Llapashtica, C., Wang, J., Kim, D.J., Xia, D., et al. (2020). TREM2 Regulates Microglial Cholesterol Metabolism upon Chronic Phagocytic Challenge. Neuron 105, 837–854.e9.

Oakley, H., Cole, S.L., Logan, S., Maus, E., Shao, P., Craft, J., Guillozet-Bongaarts, A., Ohno, M., Disterhoft, J., Van Eldik, L., et al. (2006). Intraneuronal beta-Amyloid Aggregates, Neurodegeneration, and Neuron Loss in Transgenic Mice with Five Familial Alzheimer’s Disease Mutations: Potential Factors in Amyloid Plaque Formation. J. Neurosci. 26, 10129–10140.

Ojelade, S.A., Lee, T. V., Giagtzoglou, N., Yu, L., Ugur, B., Li, Y., Duraine, L., Zuo, Z., Petyuk, V., De Jager, P.L., et al. (2019). cindr, the Drosophila Homolog of the CD2AP Alzheimer’s Disease Risk Gene, Is Required for Synaptic Transmission and Proteostasis. Cell Rep. 28, 1799–1813.e5.

Di Paolo, G., and Kim, T.-W. (2011). Linking lipids to Alzheimer’s disease: cholesterol and beyond. Nat. Rev. Neurosci. 12, 284–296.

Peña-Bautista, C., López-Cuevas, R., Cuevas, A., Baquero, M., and Cháfer-Pericás, C. (2019). Lipid peroxidation biomarkers correlation with medial temporal atrophy in early Alzheimer Disease. Neurochem. Int. 129, 104519.

Pereira, C.D., Martins, F., Wiltfang, J., Da Cruz E Silva, O.A.B., and Rebelo, S. (2017). ABC Transporters Are Key Players in Alzheimer’s Disease. J. Alzheimer’s Dis. 61, 463–485.

Quintana, A., Zanella, S., Koch, H., Kruse, S.E., Lee, D., Ramirez, J.M., and Palmiter, R.D. (2012). Fatal breathing dysfunction in a mouse model of Leigh syndrome. 122, 2359–2368.

Qureshi, Y.H., Baez, P., and Reitz, C. (2020). Endosomal Trafficking in Alzheimer’s Disease, Parkinson’s Disease, and Neuronal Ceroid Lipofuscinosis. Mol. Cell. Biol. 40, 1–12.

Ramjaun, A.R., Micheva, K.D., Bouchelet, I., and McPherson, P.S. (1997). Identification and characterization of a nerve terminal-enriched amphiphysin isoform. J. Biol. Chem. 272, 16700–16706.

Rauch, J.N., Luna, G., Guzman, E., Audouard, M., Challis, C., Sibih, Y.E., Leshuk, C., Hernandez, I., Wegmann, S., Hyman, B.T., et al. (2020). LRP1 is a master regulator of tau uptake and spread. Nature 580, 381–385.

Razzaq, A., Robinson, I.M., McMahon, H.T., Skepper, J.N., Su, Y., Zelhof, A.C., Jackson, A.P., Gay, N.J., and O’Kane, C.J. (2001). Amphiphysin is necessary for organization of the excitation-contraction coupling machinery of muscles, but not for synaptic vesicle endocytosis in Drosophila. Genes Dev. 15, 2967–2979.

Reed, T.T. (2011). Lipid peroxidation and neurodegenerative disease. Free Radic. Biol. Med. 51, 1302–1319.

Regen, F., Hellmann-Regen, J., Costantini, E., and Reale, M. (2017). Neuroinflammation and Alzheimer’s Disease: Implications for Microglial Activation. Curr. Alzheimer Res. 14.

Ristow, M., and Schmeisser, S. (2011). Extending life span by increasing oxidative stress. Free Radic. Biol. Med. 51, 327–336.

Robert, J., Button, E.B., Yuen, B., Gilmour, M., Kang, K., Bahrabadi, A., Stukas, S., Zhao, W., Kulic, I., and Wellington, C.L. (2017). Clearance of beta-amyloid is facilitated by apolipoprotein E and circulating highdensity lipoproteins in bioengineered human vessels. Elife 6, 1–24.

Rodríguez-Vázquez, M., Vaquero, D., Parra-Peralbo, E., Mejía-Morales, J.E., and Culi, J. (2015). Drosophila Lipophorin Receptors Recruit the Lipoprotein LTP to the Plasma Membrane to Mediate Lipid Uptake. PLOS Genet. 11, 1–24.

Rogaeva, E., Kawarai, T., and St. George-Hyslop, P. (2006). Genetic complexity of Alzheimer’s disease: Successes and challenges. J. Alzheimer’s Dis. 9, 381–387.

Schindelin, J., Arganda-Carreras, I., Frise, E., Kaynig, V., Longair, M., Pietzsch, T., Preibisch, S., Rueden, C., Saalfeld, S., Schmid, B., et al. (2012). Fiji: An open-source platform for biological-image analysis. Nat. Methods 9, 676–682.

Seto, E.S., Bellen, H.J., and Lloyd, T.E. (2002). When cell biology meets development: Endocytic regulation of signaling pathways. Genes Dev. 16, 1314–1336.

Sharman, M.J., Morici, M., Hone, E., Berger, T., Taddei, K., Martins, I.J., Lim, W.L.F., Singh, S., Wenk, M.R., Ghiso, J., et al. (2010). APOE genotype results in differential effects on the peripheral clearance of amyloid-β42 in APOE knock-in and knock-out mice. J. Alzheimer’s Dis. 21, 403–409.

Shen, R., Zhao, X., He, L., Ding, Y., Xu, W., Lin, S., Fang, S., Yang, W., Sung, K., Spencer, B., et al. (2020). Upregulation of RIN3 induces endosomal dysfunction in Alzheimer’s disease. Transl. Neurodegener. 9, 26–44.

Shinohara, M., Tachibana, M., Kanekiyo, T., and Bu, G. (2017). Role of LRP1 in the pathogenesis of Alzheimer’s disease: Evidence from clinical and preclinical studies. J. Lipid Res. 58, 1267–1281.

Sillitoe, R. V., Stephen, D., Lao, Z., and Joyner, A.L. (2008). Engrailed homeobox genes determine the organization of Purkinje cell sagittal stripe gene expression in the adult cerebellum. J. Neurosci. 28, 12150–12162.

Singh, A., Kukreti, R., Saso, L., and Kukreti, S. (2019). Oxidative stress: A key modulator in neurodegenerative diseases. Molecules 24, 1–20.

Steinberg, S., Stefansson, H., Jonsson, T., Johannsdottir, H., Ingason, A., Helgason, H., Sulem, P., Magnusson, O.T., Gudjonsson, S.A., Unnsteinsdottir, U., et al. (2015). Loss-of-function variants in ABCA7 confer risk of Alzheimer’s disease. Nat. Genet. 47, 445–447.

Stelzmann, R.A., Norman Schnitzlein, H., and Reed Murtagh, F. (1995). An english translation of alzheimer’s 1907 paper, “über eine eigenartige erkankung der hirnrinde.” Clin. Anat. 8, 429–431.

Stockinger, W., Sailler, B., Strasser, V., Recheis, B., Fasching, D., Kahr, L., Schneider, W.J., and Nimpf, J. (2002). The PX-domain protein SNX17 interacts with members of the LDL receptor family and modulates endocytosis of the LDL receptor. EMBO J. 21, 4259–4267.

Strittmatter, W.J., Saunders, A.M., Schmechel, D., Pericak-Vance, M., Enghild, J., Salvesen, G.S., and Roses, A.D. (1993). Apolipoprotein E: High-avidity binding to β-amyloid and increased frequency of type 4 allele in late-onset familial Alzheimer disease. Proc. Natl. Acad. Sci. U. S. A. 90, 1977–1981.

De Strooper, B., and Karran, E. (2016). The Cellular Phase of Alzheimer’s Disease. Cell 164, 603–615.

Takeda, T., Kozai, T., Yang, H., Ishikuro, D., Seyama, K., Kumagai, Y., Abe, T., Yamada, H., Uchihashi, T., Ando, T., et al. (2018). Dynamic clustering of dynamin-amphiphysin helices regulates membrane constriction and fission coupled with GTP hydrolysis. Elife 7, 1–19.

Takei, K., and Haucke, V. (2001). Clathrin-mediated endocytosis: Membrane factors pull the trigger. Trends Cell Biol. 11, 385–391.

Tarling, E.J., Vallim, T.Q. d. A., and Edwards, P.A. (2013). Role of ABC transporters in lipid transport and human disease. Trends Endocrinol. Metab. 24, 342–350.

Teresa, J.C., Fernado, C., Nancy, M.R., Gilberto, V.A., Alberto, C.R., and Roberto, R.R. (2020). Association of genetic variants of ABCA1 with susceptibility to dementia: (SADEM study). Metab. Brain Dis. 35, 915–922.

Thapa, A., and Carroll, N.J. (2017). Dietary modulation of oxidative stress in Alzheimer’s disease. Int. J. Mol. Sci. 18, 1583–1596.

Tönnies, E., and Trushina, E. (2017). Oxidative Stress, Synaptic Dysfunction, and Alzheimer’s Disease. J. Alzheimer’s Dis. 57, 1105–1121.

Turton, J., and Morgan, K. (2013). ATP-binding cassette, subfamily A (ABC1), member 7 (ABCA7). In Genetic Variants in Alzheimer’s Disease, (Springer New York), pp. 135–158.

Ubelmann, F., Burrinha, T., Salavessa, L., Gomes, R., Ferreira, C., Moreno, N., and Guimas Almeida, C. (2017). Bin1 and CD 2 AP polarise the endocytic generation of beta-amyloid. EMBO Rep. 18, 102–122.

Venken, K.J.T., He, Y., Hoskins, R.A., and Bellen, H.J. (2006). P[acman]: A BAC transgenic platform for targeted insertion of large DNA fragments in D. melanogaster. Science (80-.). 314, 1747–1751.

Verghese, P.B., Castellano, J.M., Garai, K., Wang, Y., Jiang, H., Shah, A., Bu, G., Frieden, C., and Holtzman, D.M. (2013). ApoE influences amyloid-β (Aβ) clearance despite minimal apoE/Aβ association in physiological conditions. Proc. Natl. Acad. Sci. U. S. A. 110, E1807–E1816.

Verstreken, P., Koh, T.W., Schulze, K.L., Zhai, R.G., Hiesinger, P.R., Zhou, Y., Mehta, S.Q., Cao, Y., Roos, J., and Bellen, H.J. (2003). Synaptojanin is recruited by endophilin to promote synaptic vesicle uncoating. Neuron 40, 733–748.

Vina, J., LLoret, A., Giraldo, E., C. Badia, M., and D. Alonso, M. (2011). Antioxidant Pathways in Alzheimers Disease: Possibilities of Intervention. Curr. Pharm. Des. 17, 3861–3864.

Wahrle, S.E., Jiang, H., Parsadanian, M., Legleiter, J., Han, X., Fryer, J.D., Kowalewski, T., and Holtzman, D.M. (2004). ABCA1 is required for normal central nervous system apoE levels and for lipidation of astrocyte-secreted apoE. J. Biol. Chem. 279, 40987–40993.

Wahrle, S.E., Jiang, H., Parsadanian, M., Kim, J., Li, A., Knoten, A., Jain, S., Hirsch-Reinshagen, V., Wellington, C.L., Bales, K.R., et al. (2008). Overexpression of ABCA1 reduces amyloid deposition in the PDAPP mouse model of Alzheimer disease. J. Clin. Invest. 118, 671–682.

Wang, Y., and Mandelkow, E. (2016). Tau in physiology and pathology. Nat. Rev. Neurosci. 17, 5–21.

Wang, S., Tan, K.L., Agosto, M.A., Xiong, B., Yamamoto, S., Sandoval, H., Jaiswal, M., Bayat, V., Zhang, K., Charng, W.L., et al. (2014). The Retromer Complex Is Required for Rhodopsin Recycling and Its Loss Leads to Photoreceptor Degeneration. PLoS Biol. 12, 1–20.

Wang, Y., Cella, M., Mallinson, K., Ulrich, J.D., Young, K.L., Robinette, M.L., Gilfillan, S., Krishnan, G.M., Sudhakar, S., Zinselmeyer, B.H., et al. (2015). TREM2 lipid sensing sustains the microglial response in an Alzheimer’s disease model. Cell 160, 1061–1071.

Ward, A., Crean, S., Mercaldi, C.J., Collins, J.M., Boyd, D., Cook, M.N., and Arrighi, H.M. (2012). Prevalence of Apolipoprotein E4 Genotype and Homozygotes (APOE e4/4) among Patients Diagnosed with Alzheimer’s Disease: A Systematic Review and Meta-Analysis. Neuroepidemiology 38, 1–17.

Wellington, C. (2004). Cholesterol at the crossroads: Alzheimer’s disease and lipid metabolism. Clin. Genet. 66, 1–16.

Wojtunik-Kulesza, K.A., Oniszczuk, A., Oniszczuk, T., and Waksmundzka-Hajnos, M. (2016). The influence of common free radicals and antioxidants on development of Alzheimer’s Disease. Biomed. Pharmacother. 78, 39–49.

Wong, M.W., Braidy, N., Poljak, A., Pickford, R., Thambisetty, M., and Sachdev, P.S. (2017). Dysregulation of lipids in Alzheimer’s disease and their role as potential biomarkers. Alzheimer’s Dement. 13, 810–827.

Zabel, M., Nackenoff, A., Kirsch, W.M., Harrison, F.E., Perry, G., and Schrag, M. (2018). Markers of oxidative damage to lipids, nucleic acids and proteins and antioxidant enzymes activities in Alzheimer’s disease brain: A meta-analysis in human pathological specimens. Free Radic. Biol. Med. 115, 351–360.

Zelhof, A.C., Bao, H., Hardy, R.W., Razzaq, A., Zhang, B., and Doe, C.Q. (2001). Drosophila Amphiphysin is implicated in protein localization and membrane morphogenesis but not in synaptic vesicle endocytosis. Development 128, 5005–5015.

Zhang, C., and Liu, P. (2017). The lipid droplet: A conserved cellular organelle. Protein Cell 1–5.

Zhang, B., Koh, Y.H., Beckstead, R.B., Budnik, V., Ganetzky, B., and Bellen, H.J. (1998). Synaptic vesicle size and number are regulated by a clathrin adaptor protein required for endocytosis. Neuron 21, 1465–1475.

Zhang, H., Huang, T., Hong, Y., Yang, W., Zhang, X., Luo, H., Xu, H., and Wang, X. (2018). The retromer complex and sorting nexins in neurodegenerative diseases. Front. Aging Neurosci. 10, 1–11.

Zhang, Y., Chen, K., Sloan, S.A., Bennett, M.L., Scholze, A.R., O’Keeffe, S., Phatnani, H.P., Guarnieri, P., Caneda, C., Ruderisch, N., et al. (2014). An RNA-sequencing transcriptome and splicing database of glia, neurons, and vascular cells of the cerebral cortex. J. Neurosci. 34, 11929–11947.

Zhang, Y., Sloan, S.A., Clarke, L.E., Caneda, C., Plaza, C.A., Blumenthal, P.D., Vogel, H., Steinberg, G.K., Edwards, M.S.B.B., Li, G., et al. (2016). Purification and Characterization of Progenitor and Mature Human Astrocytes Reveals Transcriptional and Functional Differences with Mouse Highlights. Neuron 89, 37–53.

Zhu, X.C., Tan, L., Wang, H.F., Jiang, T., Cao, L., Wang, C., Wang, J., Tan, C.C., Meng, X.F., and Yu, J.T. (2015). Rate of early onset Alzheimer’s disease: A systematic review and meta-analysis. Ann. Transl. Med. 3, 38–43.

